# An EYA3/NF-κB/CCL2 signaling axis suppresses cytotoxic NK cells in the pre-metastatic niche to promote triple negative breast cancer metastasis

**DOI:** 10.1101/2024.07.31.606072

**Authors:** Sheera R. Rosenbaum, Connor J. Hughes, Kaiah M. Fields, Stephen Connor Purdy, Annika Gustafson, Arthur Wolin, Drake Hampton, Nicholas Turner, Christopher Ebmeier, James C. Costello, Heide L. Ford

## Abstract

Patients with Triple Negative Breast Cancer (TNBC) exhibit high rates of metastases and poor prognoses. The Eyes absent (EYA) family of proteins are developmental transcriptional cofactors/phosphatases that are re-expressed and/or upregulated in numerous cancers. Herein, we demonstrate that EYA3 correlates with decreased survival in breast cancer, and that it strongly, and specifically, regulates metastasis via a novel mechanism that involves NF-kB signaling and an altered innate immune profile at the pre-metastatic niche (PMN).

Remarkably, restoration of NF-kB signaling downstream of *Eya3* knockdown (KD) restores metastasis *without* restoring primary tumor growth, isolating EYA3/NF-kB effects to the metastatic site. We show that secreted CCL2, regulated downstream of EYA3/NF-kB, specifically decreases cytotoxic NK cells in the PMN and that re-expression of *Ccl2* in *Eya3*-KD cells is sufficient to rescue activation/levels of cytotoxic NK cells *in vitro and* at the PMN, where EYA3-mediated decreases in cytotoxic NK cells are required for metastatic outgrowth.

Importantly, analysis of public breast cancer datasets uncovers a significant correlation of EYA3 with NF-kB/CCL2, underscoring the relevance of EYA3/NF-kB/CCL2 to human disease. Our findings suggest that inhibition of EYA3 could be a powerful means to re-activate the innate immune response at the PMN, inhibiting TNBC metastasis.

**Significance:** EYA3 promotes metastasis of TNBC cells by promoting NF-kB-mediated CCL2 expression and inhibiting cytotoxic NK cells at the pre-metastatic niche, highlighting a potential therapeutic target in this subset of breast cancer.

## Introduction

Triple-negative breast cancer (TNBC), marked by low/absent expression of estrogen receptor (ER), progesterone receptor (PR), and the human epidermal growth factor receptor 2 (HER2)^1–3^, represents 12-20% of all breast cancers^2,4,5^. TNBC has a predilection for women under the age of 40^2^, and is a molecularly diverse disease that is often highly aggressive in nature^1,2,6^. Due to the lack of targetable receptor expression, treatment is often limited to conventional surgery, radiation, and chemotherapeutic approaches^7^, leading to poor overall survival for metastatic TNBC patients relative to other subtypes of breast cancer^2^. With this in mind, further exploration and development of alternative treatment options such as antibody-drug conjugates, PARP inhibitors, or immune checkpoint inhibitors suggest targeted approaches may provide additional benefit in specific cases^6^. Unfortunately, stage IV TNBC remains nearly uniformly lethal^8^. Thus, a greater understanding of the metastatic processes utilized by TNBC cells is essential to improve outcomes in the future.

Embryonic programs are often re-activated in cancer. The vertebrate Eyes absent protein family, consisting of EYAs1-4, are critical for the development of multiple organ systems in mammals, including the kidney^9–12^, ear^10,11,13,14^, muscle^12,15^, and craniofacial structures^10,11,15^. The developmental functions of EYAs are conferred through three distinct activities: 1) co-activation of transcription, mediated largely via an interaction with Six family homeoproteins^16–18^, 2) Tyrosine (Tyr) phosphatase activity, mediated through an intrinsic haloacid dehalogenase (HAD) Tyr phosphatase domain in the C-terminal Eya domain (ED)^12,16,19^, and 3) Serine/Threonine (Ser/Thr) phosphatase activity found in the N-Terminal domain (ED2)^12^, and mediated via an interaction with the B55α subunit of PP2A^20,21^. While expression of EYA proteins is generally low or absent after development is complete^22^, increased- or re-expression of EYAs is observed in many cancers, including breast^20,23–25^, lung^26,27^, ovarian^28,29^, and malignant peripheral nerve sheath tumors^30^, amongst others. In the context of development and cancer, EYAs promote many critical cellular processes, including epithelial-to-mesenchymal transition^24^, cell cycle progression and proliferation^23,31^, DNA repair and cell survival^30,32–34^, and cellular migration, invasion^23^, and angiogenesis^35^. Through an interaction with PP2A-B55α, EYAs stabilize the proto-oncogene c-MYC^20^, facilitating tumor progression^21^. We previously showed that EYA’s regulation of MYC promotes PD-L1 expression in TNBC, leading to exhaustion of cytotoxic CD8+ T-cells and enhanced primary tumor growth^25^. Of interest, EYAs have also been shown to regulate innate immune signaling in response to cytoplasmic DNA/RNA in both Drosophila^36^ and mouse embryonic fibroblasts^37^, suggesting that these functions may be conserved in the context of cancer. However, a role for EYA3 in the regulation of innate immune responses to tumors has never been demonstrated.

The NF-kB family of transcription factors drives the expression of diverse target genes, including cytokines/chemokines, cell-surface receptors, pro-survival genes, stress response genes, and growth factors^38–41^. The canonical NF-kB signaling pathway, which signals through the IkB Kinase (IKK) complex^42^, is activated through phosphorylation of serines 177 and 181 on IKKβ^43^, leading to phosphorylation of IκB ^44,45^. IκB phosphorylation results in its degradation and subsequent release of RELA/p50 for nuclear translocation. Once nuclear, RELA/p50 activates numerous genes that regulate growth, survival, innate immunity and inflammation^44^. While NF- kB can be anti-tumorigenic^46^, it is often pro-tumorigenic and metastatic when activated in cancer cells, through altering transcription of genes that lead to increases in tumor cell proliferation, survival, invasion, therapy resistance, inflammation, and changes in both adaptive and innate immune responses to tumors^46–50^. These many functions of NF-kB initially made it an attractive cancer target, but unfortunately, its inhibition is associated with significant toxicities^1,47,51,52^.

During metastatic progression, cancer cells prime distant sites for metastasis, establishing a pre-metastatic niche that is favorable for cancer cell seeding^53–55^. Formation of the pre-metastatic niche involves suppression of anti-tumor immune responses at secondary sites prior to the arrival of metastatic cells, requiring a complex crosstalk between tumor-derived factors, stromal cells, and immune cells^56–61^. Numerous cytokines influence immune cells in the pre-metastatic niche^62^, and CCL2, which can be regulated by NF-kB^63^, has been implicated in promoting the recruitment of both pro-metastatic inflammatory monocytes^59^ and anti-metastatic neutrophils^64^ to the lung. While many immune cells regulate the pre-metastatic niche, cytotoxic NK cells are critical for the elimination of metastatic cells in the lung, as NK cell depletion results in increased metastatic outgrowth in mouse models of breast and lung cancers^65–67^.

Accumulating evidence indicates that NK cell activity is associated with a reduced risk of metastasis, and that NK cells can be exploited therapeutically to treat metastatic disease^68^. However, NK cells can also be reeducated by cancer cells to promote tumor growth and metastasis^69,70^. Thus, interactions in the immune pre-metastatic niche are complex, and a comprehensive understanding of the interaction between cancer cells and immune cells during the pre-metastatic window is lacking.

In this study, we demonstrate that EYA3 promotes NF-kB signaling in TNBC by enhancing nuclear localization of the RELA transcription factor, and that NF-kB signaling downstream of Eya3 is necessary for spontaneous metastasis of murine TNBC cells, but not for effects on primary tumor growth. We show that EYA3-mediated NF-kB signaling results in increased CCL2 expression, which limits cytotoxic NK cells in the pre-metastatic niche and inhibits cytotoxic NK cell activation potential *in vitro*. Finally, we demonstrate that this EYA3/NF- kB/CCL2 axis extends to human breast cancer, as these genes and signaling pathways strongly correlate in patient datasets. There remains a dire need for new therapeutic strategies to treat patients with metastatic TNBC. Thus, these findings suggest a possible future avenue for inhibiting EYA3 as a means to target both MYC and NF-kB signaling, thus inhibiting both primary and metastatic TNBC^27,71–78^.

## Results

### EYA3 promotes spontaneous metastasis of murine TNBC cells

We previously demonstrated that EYA3 promotes primary tumor growth in breast cancer (BC) via regulating a MYC/PDL1 axis and suppressing the adaptive immune response to the tumor^25^. However, whether EYA3 may also regulate spontaneous metastasis, which is the leading cause of BC related deaths^79^, and thus significantly affects the prognosis of women with BC, remained unknown. To address this question, we first analyzed overall survival (OS) in the METABRIC patient dataset for patients with high versus low *EYA3*^80–82^, and found that patients with high *EYA3* expression (top quartile) exhibited decreased survival compared to those with low *EYA3* expression (bottom quartile) (Fig. 1A). Given the association of *EYA3* with worsened survival, and that previous studies have only evaluated the effect of EYA3 on primary tumor growth^25^ and late-stage metastasis (the latter in immune compromised settings)^21^, we asked whether EYA3 could promote the entire metastatic cascade from the primary tumor in immune-competent models of TNBC. We focused specifically on TNBC, given that EYA3 is more highly expressed in this subset^25^ and there is a dire need for new therapies in this breast cancer subgroup. We performed knockdown (KD) of *Eya3* using two independent shRNAs in the 66cl4 murine TNBC cell line^83^ (Figs. 1B-C) after which we injected luciferase-tagged 66cl4 shRNA scramble control (SCR) and *Eya3* KD cells into the mammary fat pad of immune-competent syngeneic BALB/c mice (experimental layout, Fig. 1D). Primary tumor growth was tracked over time by caliper measurement and, in line with previous findings^25^, we observed a strong reduction in primary tumor growth with *Eya3* KD (Fig. 1E). In our first experiment, we removed the primary tumor when the largest tumor reached 1cm^3^ and continued to image the mice weekly after tumor removal using the IVIS bioluminescent imager to detect metastases. We observed a strong decrease in bioluminescence in the lung window for mice injected with 66cl4 *Eya3* KD cells compared to SCR cells (Fig. 1F, Suppl. Fig. 1A). As the 66cl4 SCR primary tumors were much larger than the *Eya3* KD tumors at the time of removal (Fig. 1E), and as tumor size is known to correlate with incidence of distant metastases in TNBC^84^, we further asked whether the increased tumor burden observed in the lungs of 66cl4 SCR mice after primary tumor removal was simply a consequence of increased overall tumor burden, or whether EYA3 was directly regulating a pro-metastatic phenotype. We thus repeated the metastasis experiment, but instead removed each tumor at the same size (1cm^3^) rather than at the same time. After primary tumor removal, the mice were again imaged weekly and bioluminescence intensity was averaged across each group and plotted based on the number of days after tumor removal.

**Figure 1:**
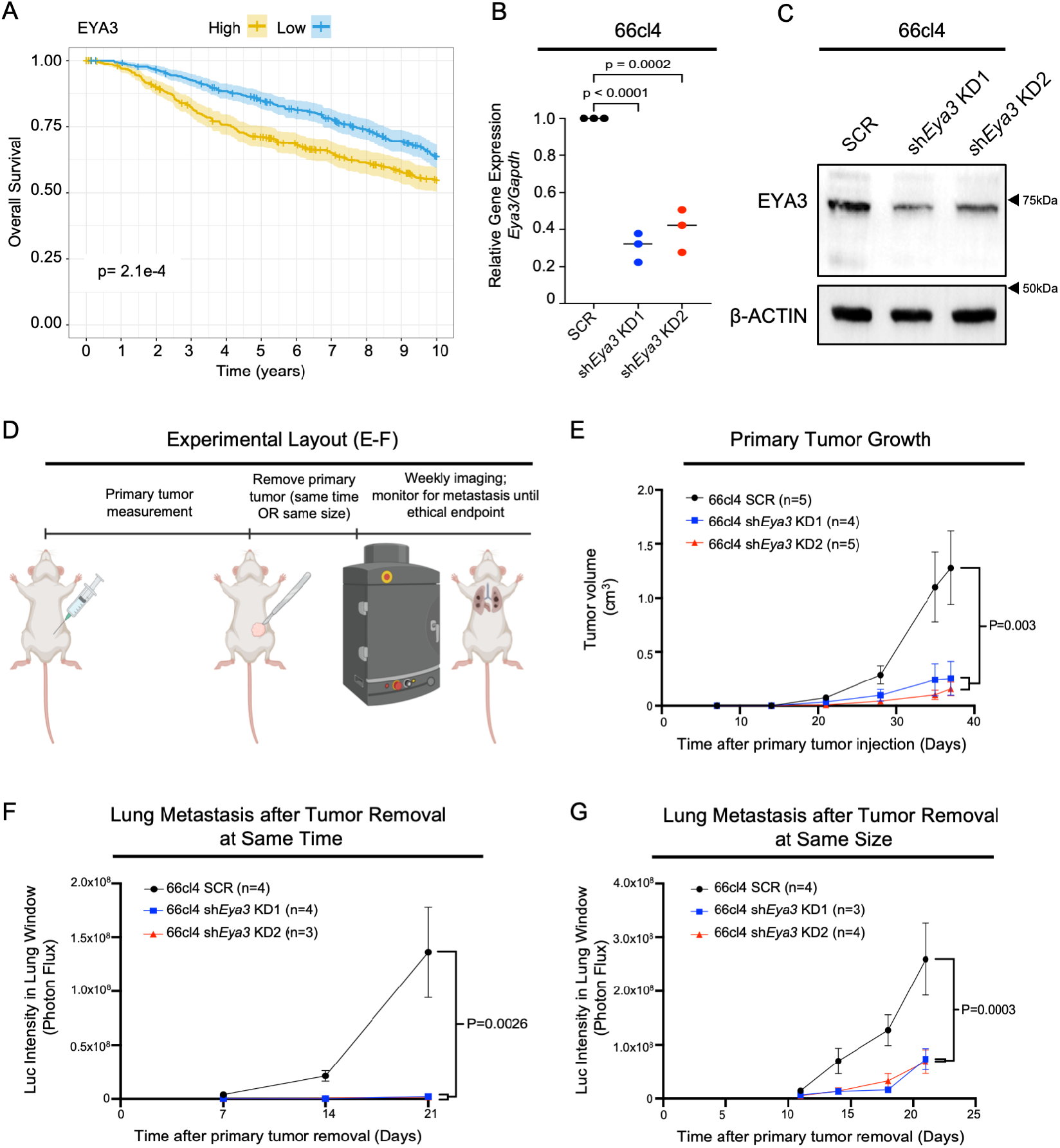
EYA3 correlates with poor prognosis in human TNBC, and promotes primary tumor growth and metastasis in immune-competent models of TNBC. (A) Gene expression for *EYA3* was used to stratify the overall cohort into quartiles, and the top and bottom quartiles were compared for overall survival using the survival and survminer R packages. Overall survival was analyzed with patients living past 10 years right censored; the log-rank test was used to calculate statistical significance. (B) *Eya3* mRNA expression, as measured using real time qRT-PCR. *Eya3* expression was normalized to *Gapdh* levels. (C) Western blot analysis of EYA3 in 66cl4 SCR and *Eya3* KD cells. (D) Experimental design for the *in vivo* experiment with data in panels E-F (largest tumor reaches 1cm^3^) and G (size-matched tumors, 1cm^3^). (E) Primary tumor growth via caliper measurement after orthotopic injection of luciferase-tagged 66cl4 SCR, sh*Eya3* KD1, and sh*Eya3* KD2 cells into BALB/c mice. (F) Bioluminescence intensity in the lung window over time after primary tumor removal for mice injected with 66cl4 SCR, sh*Eya3* KD1, and sh*Eya3* KD2. Two mice were euthanized and excluded due to surgical complications, and one (KD2) mouse was excluded from analysis after calculated to be a significant outlier via Grubbs’ test. (G) Bioluminescence intensity in lung window of mice injected with 66cl4 SCR, sh*Eya3* KD1, or sh*Eya3* KD2 cells after orthotopic injection into the mammary fat pad followed by primary tumor removal when all tumors reached the size of 1cm^3^. Statistical analysis for E, F, and G was performed using longitudinal mixed model analysis in R. Mean±SEM shown.

Data from this second study revealed that EYA3 plays a distinct role in metastasis of TNBC, as a significant decrease in bioluminescence was still observed in the *Eya3* KD vs SCR cells when tumors were removed at the same size (Fig. 1G, Suppl. Fig. 1B).

To determine whether this finding holds true in a second immune-competent model of TNBC, and to further confirm that differences in primary tumor size are not contributing to EYA3 effects on metastasis, we knocked down *Eya3* in E0771 cells (Suppl. Fig. 1C, 1D), which are also considered TNBC^85^, though this designation is controversial^86^. We injected E0771 murine breast cancer cells with or without *Eya3* KD into the tail vein of syngeneic C57BL/6 mice (thus removing effects of primary tumor size, but maintaining an immune competent environment).

After 21 days we observed that several mice reached a humane endpoint without showing any signs of luciferase signal, leading us to speculate that the luciferase tag had been lost in these cells, as luciferase tagging has been shown to promote increased immune clearance in other murine breast cancer models^87^. All E0771 cells express a puromycin resistance gene as a part of the shRNA plasmid; thus, we used this marker to identify tumor cells in the lungs of injected mice. We sacrificed all mice at 23 days post-injection, isolated RNA from the lungs, and performed qRT-PCR on these samples compared to RNA isolated from lung tissue from a tumor-free mouse (used as a negative control). In line with our findings in the 66cl4 model, we detected reduced expression of the puromycin resistance gene in the lungs of mice that were injected with E0771 *Eya3* KD cells when compared to the SCR control cells (Suppl. Fig. 1E).

Thus, our data demonstrate that in two immune-competent models, EYA3 mediates metastasis to the lungs.

### EYA3 promotes innate immune gene expression and NF-kB activation in murine TNBC cells

To determine how EYA3 may be promoting spontaneous metastasis, we performed bulk RNA-sequencing (RNA-seq) on 66cl4 SCR and *Eya3* KD cells. Sh*Eya3* KD1 cells had a higher degree of *Eya3* KD than sh*Eya3* KD2 cells (Fig. 1B, Suppl. Fig. 2A), though *Eya3* was knocked down in both groups. Gene set enrichment analysis (GSEA) revealed that several of the top 15 downregulated gene sets upon *Eya3* KD relate to innate immune signaling (Fig. 2A), including the Hallmark Inflammatory Response (Fig. 2A, Suppl. Fig. 2B) and Hallmark TNFɑ Signaling via NF-kB (Fig. 2A, 2B) gene sets. The heat map of TNFɑ Signaling via NF-kB genes demonstrates a dose-dependent change in expression of most genes in the pathway with *Eya3* KD (Suppl. Fig. 2C), where a larger degree of *Eya3* KD (sh*Eya3* KD1) correlates with a greater change in the expression of genes in this geneset. Of note, MYC targets were also observed in our GSEA analysis (Fig. 2A and Suppl. Fig. 2D, 2E), corroborating previous findings that *Myc* mRNA and protein is regulated by EYA3 in TNBC^21,25^. Further proteomic analysis confirmed that the top gene sets reduced in sh*Eya3* cells include inflammatory and MYC signaling gene sets (Suppl. Fig. 2F, G). These data demonstrate that in addition to regulating MYC, as previously reported^21,25^, EYA3 controls another critical tumor promotional pathway in TNBC: NF- kB/inflammatory signaling. Because EYA3 has not previously been shown to control NF-kB/ inflammatory signaling in TNBC, and because NF-kB has been implicated in metastasis^88–92^, we decided to explore this finding further.

**Figure 2:**
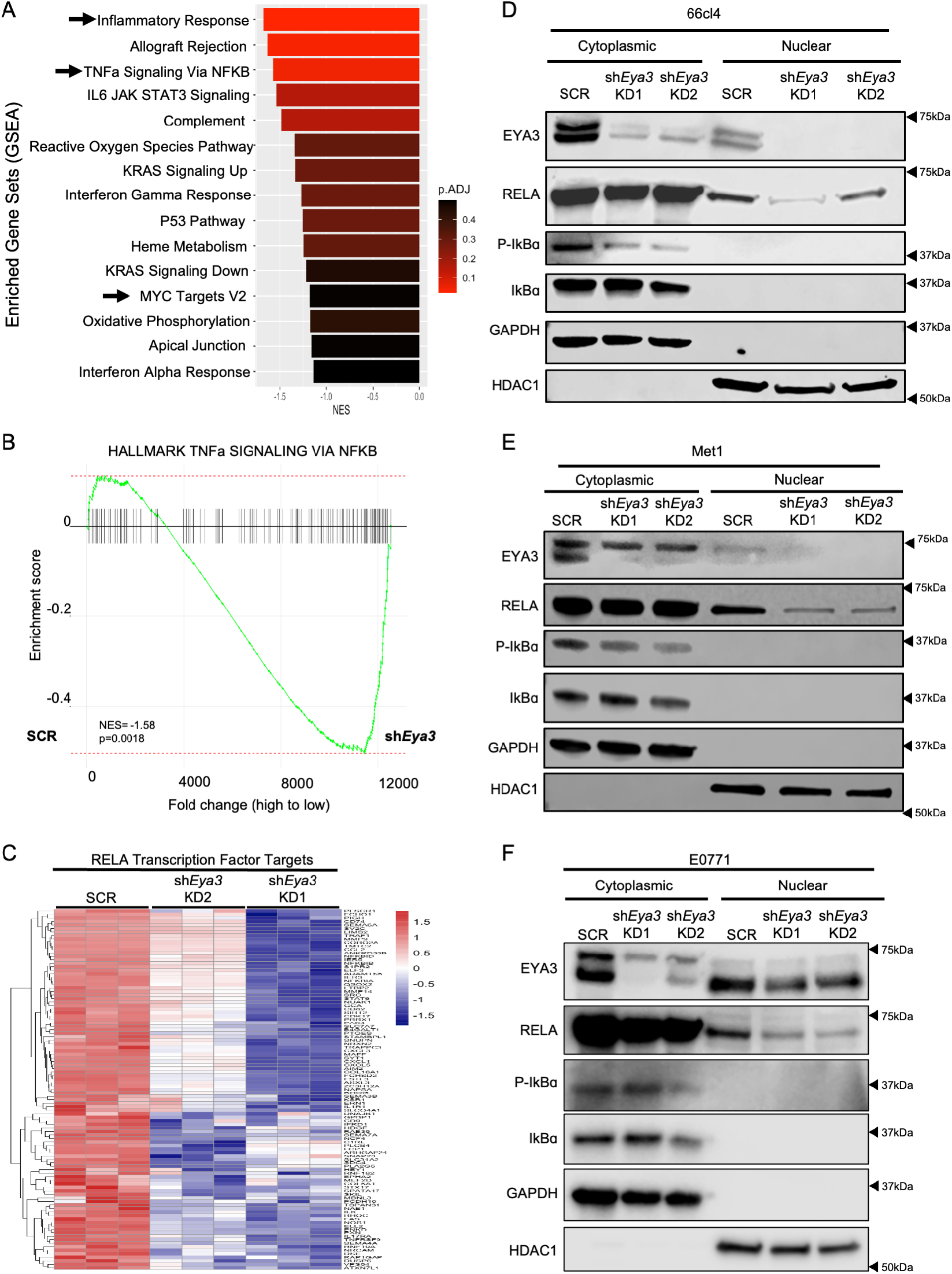
EYA3 promotes NF-kB signaling in TNBC cells. (A) RNA-seq data was analyzed for top pathways enriched in the SCR group compared to the combined *Eya3* KD groups. Shown are the top ten enriched Hallmark gene sets by GSEA, plotted in order of normalized enrichment score. (B) GSEA plot of Hallmark TNFa Signaling via NF-kB gene set between 66cl4 SCR and combined *Eya3* KD cells. (C) Heat map of RELA transcription factor target genes between 66cl4 SCR and *Eya3* KD cells. (D-F) Western blot analyses of cytoplasmic and nuclear protein fractions of 66cl4, Met1, and E0771 SCR and *Eya3* KD cells for markers of NF-kB activation.

We thus first asked whether EYA3 may be regulating a single transcription factor within the NF-kB pathway that is responsible for the alterations to the inflammatory signatures. To answer this question, we performed transcription factor enrichment analysis to determine whether genes altered by *Eya3* KD were significantly enriched within the target genes of a single transcription factor. We found that genes regulated by *Eya3* were significantly enriched within target genes of the RELA transcription factor (TF), which accounted for 9/10 of the top enriched TF motifs (with the remaining 1/10 being another NF-kB transcription factor, c-Rel or Rel) (Suppl. Fig. 2H). RELA is the canonical TF for the NF-kB pathway. Thus, alterations to genes that are bound by RELA would be expected to lead to changes in the Hallmark TNFɑ Signaling via NF-kB pathway (Fig. 2B). Heat maps of RELA target genes confirmed this finding, with dose-dependent changes observed in the expression of many RELA target genes in the *Eya3* KD1 vs. KD2 cells, when compared to the SCR cells (Fig. 2C). These findings strongly suggest that EYA3 is regulating RELA transcriptional activity in murine TNBC cells.

To determine the mechanism by which EYA3 is regulating RELA target genes, we first examined whether *Eya3* KD alters RELA protein levels. In the 66cl4 cell line and two additional murine TNBC cell lines, Met1^93^ and E0771, we observed that *Eya3* KD led to modest reductions in total RELA protein levels (Suppl. Fig. 2I, J, K). Because RELA protein is expressed in an inactive state in the cytoplasm, and can only activate downstream gene transcription after translocating to the nucleus^94^, we further examined nuclear localization of RELA as a readout for NF-kB pathway activation. To this end, we performed fractionation of nuclear and cytoplasmic proteins in the 66cl4, Met1, and E0771 SCR and *Eya3* KD cells and found that while cytoplasmic levels of RELA were unchanged, nuclear RELA levels were reduced with *Eya3* KD in all three contexts (Figs. 2D, 2E, 2F). Furthermore, we observed decreased phosphorylation of IκBα (Fig. 2D, 2E, 2F), an event that occurs upstream of RELA nuclear localization. Using an NF-kB promoter luciferase reporter in the Met1 cell line^95^, we further corroborated our finding that *Eya3* KD decreases NF-kB mediated transcription in a second TNBC system (Suppl. Fig. 2L). Taken together, these data demonstrate that EYA3 promotes nuclear translocation of RELA, leading to an induction of RELA-mediated transcription in murine TNBC cells.

### EYA3-mediated NF-kB signaling promotes metastasis but not primary tumor growth in an immune-competent mouse model

To determine whether EYA3-mediated NF-kB activation is responsible for the pro-metastatic phenotype of EYA3, we transduced 66cl4, Met1, and E0771 SCR and *Eya3* KD cells with either an empty vector plasmid (EV) or a plasmid driving expression of a phospho-mimetic form of the NF-kB activator kinase IKK2 (also referred to as IKKβ), which signals upstream of IκBα in the NF-kB pathway. The mutant protein (IKK2-SE), which has phospho-mimetic substitutions (S->E) of the activating residues Ser177 and Ser181, drives constitutively active NF-kB pathway activation^43^. We were unable to select Met1 *Eya3* KD lines expressing this construct (likely due to growth arrest and apoptotic effects of constitutively active NF-kB signaling, which have previously been reported^96,97^). However, we were able to select both 66cl4 and E0771 *Eya3* KD lines in which the *IKK2-SE* plasmid was introduced. Western blot (WB) analysis of fractionated cytoplasmic and nuclear protein in the 66cl4 and E0771 cell lines demonstrated that expression of IKK2-SE rescues nuclear RELA levels and IκBα phosphorylation in 66cl4 and E0771 *Eya3* KD cells (Fig. 3A, Suppl. Fig. 3A). In addition, RNA-sequencing showed a strong enrichment for the Hallmark NF-kB Signaling via TNFα and Hallmark Inflammatory Response gene sets with *IKK2-SE* addback in 66cl4 *Eya3* KD cells (Fig. 3B, 3C); a finding that was confirmed by performing qRT-PCR of NF-kB target genes in E0771 cells with *Eya3* KD +/- *IKK2-SE* addback (Suppl. Fig. 3B). Together, these data demonstrate that expression of IKK2-SE is sufficient to increase NF-kB activation in the context of *Eya3* KD. Importantly, while we have previously shown that EYA3 regulates MYC^21,25^, MYC signatures were not rescued by restoration of NF-kB signaling as Hallmark MYC gene sets were not top enriched hits when comparing *Eya3* KD2+*IKK2-SE* to *Eya3* KD2+EV (Suppl Fig. 3C), demonstrating that NF-kB signaling downstream of EYA3 is not responsible for EYA3-mediated activation of MYC. In line with this finding, *IKK2-SE* expression did not rescue the effects of EYA3 on proliferation (Suppl. Fig 4A, 4B), which may thus be mediated, at least in part, by MYC downstream of EYA3. In this way, we were able to separate the EYA3/NFkB axis from the Eya3/MYC axis for subsequent studies.

**Figure 3:**
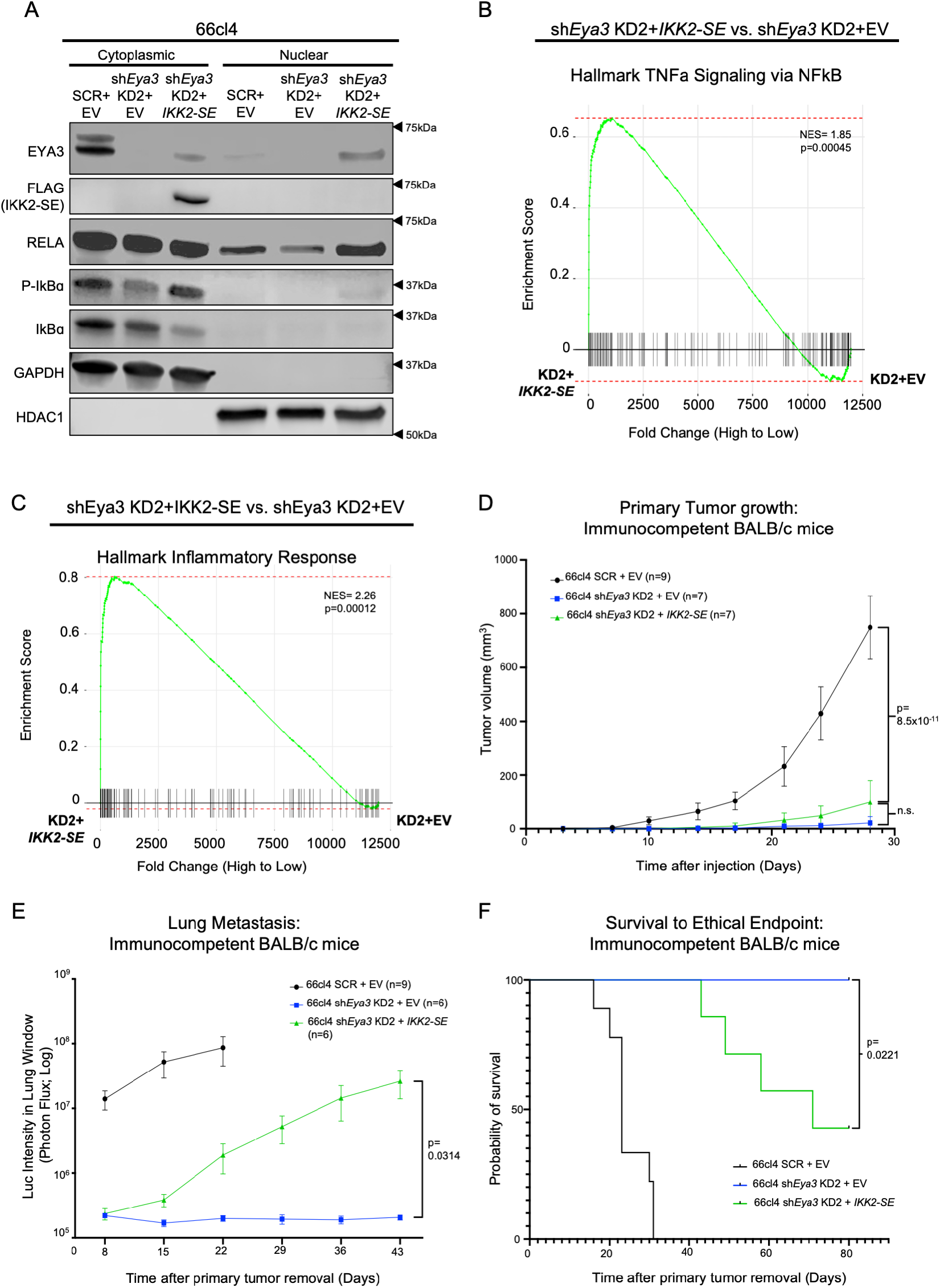
EYA3-mediated NF-kB signaling promotes metastasis and worsens survival in immune- competent mouse models of TNBC. (A) Western blot analysis of cytoplasmic and nuclear protein fractions of 66cl4 SCR+EV, sh*Eya3* KD2+EV and sh*Eya3* KD2+*IKK2-SE* for Flag-IKK2-SE and markers of NF-kB activation. (B) GSEA plot of Hallmark TNFa Signaling via NFkB gene set from RNA-seq analysis comparing gene expression in 66cl4 sh*Eya3* KD2+*IKK2-SE* versus sh*Eya3* KD2+EV cells. (C) GSEA plot of Hallmark Inflammatory Response gene set from RNA-seq analyzed in B. (D) Primary tumor growth, as measured by caliper measurement, of 66cl4 SCR+EV, sh*Eya3* KD2+EV, and sh*Eya3* KD2+*IKK-SE* cells after orthotopic injection into the mammary fat pad of BALB/c mice. (E) Bioluminescence intensity in lung window of BALB/c mice after primary tumor removal for BALB/c mice injected with 66cl4 SCR+EV, sh*Eya3* KD2+EV and sh*Eya3* KD2+*IKK2-SE* cells. Statistical analysis for D-E performed using a longitudinal mixed model in R. (F) Overall survival to an ethical endpoint for mice from E. Statistical analysis was performed using log-rank test to compare survival.

To determine whether EYA3 mediates metastasis via regulating NF-kB signaling, we injected 66cl4 SCR, *Eya3* KD, and *Eya3* KD+ *IKK2SE* cells into the orthotopic site of immune-competent syngeneic BALB/c mice. Primary tumors were then removed (at day 36) and the mice were monitored over time for evidence of lung metastasis using bioluminescent imaging. Surprisingly, NF-kB reactivation in *Eya3* KD cells did ***not*** rescue primary tumor growth in immune-competent mice (Fig. 3D), but it ***did*** significantly rescue lung metastasis (Fig. 3E, Suppl. Fig. 4C). In line with the increased metastasis, restoration of NF-kB signaling downstream of *Eya3* KD also significantly reduced survival (Fig. 3F).

Given that NF-kB has been previously implicated in cell autonomous phenotypes linked to metastasis, such as migration^91^, we asked whether *IKK2-SE* expression, downstream of *Eya3* KD, alters migration *in vitro*, and metastasis *in vivo* in an immune-deficient NOD/SCIDγ (NSG) model. Indeed, IKK2-SE rescued EYA3-mediated migration defects *in vitro* (Suppl. Fig. 5A-5D), and metastatic burden in NSG mice (Suppl. Fig. 5E-5H), demonstrating that tumor cell-autonomous effects contribute to EYA3/NF-kB-mediated metastasis. However, given the profound effects on metastasis observed in the immune competent model (Fig. 3E), and restoration of inflammatory signaling upon NF-kB reactivation in *Eya3* KD cells (Fig. 3C), we focused our attention moving forward on determining whether EYA3 may also be altering the immune TME to promote metastasis.

### TNBC-expressed EYA3 remodels the pre-metastatic lung immune microenvironment

We previously reported that *Eya3* KD significantly alters multiple adaptive and innate immune populations within the primary tumor^25^, and RNA-seq analysis of 66cl4 SCR versus *Eya3* KD cells shows alterations in inflammatory signaling pathways, including multiple cytokines and cytokine receptors (Fig. 2A, Suppl. Fig. 2B). These data led us to hypothesize that EYA3 may be changing the TME not only in the primary tumor but also at the PMN, contributing to the large differences in metastasis observed with *Eya3* KD in the immune competent setting. As cancers progress to the metastatic stage, cells from the primary tumor prepare distant sites for colonization, in part, by altering the immune microenvironment in the pre-metastatic niche^98^. To determine whether EYA3 affects immune cells in the pre-metastatic niche, we injected 66cl4 cells with or without *Eya3* KD into the mammary fat pad of syngeneic mice and allowed palpable tumors (>∼50mm^3^) to form (Fig. 4A). In line with previous experiments (Fig. 1E), cells with *Eya3* KD grew at a slower rate at the orthotopic site (Fig. 4B), and took longer to form palpable tumors (Fig. 4C). Once tumors were palpable, mice were sacrificed and lungs were processed into a single cell suspension. A small portion of this single cell suspension was used to isolate RNA, which was then probed for expression of luciferase mRNA by qRT-PCR, to determine whether any cancer cells were detectable in the lungs at this early timepoint. Our assay was sensitive enough to detect 1 cancer cell in 10^5^ lung cells (Suppl. Fig. 6A), and with this level of sensitivity, we did not detect any cancer cells in the lungs of mice with 50mm^3^ palpable primary orthotopic tumors (Suppl. Fig. 6B). Thus, our data suggest that our experimental conditions could be used to assess the “pre-metastatic” phase before micro or macro metastases are detected.

**Figure 4:**
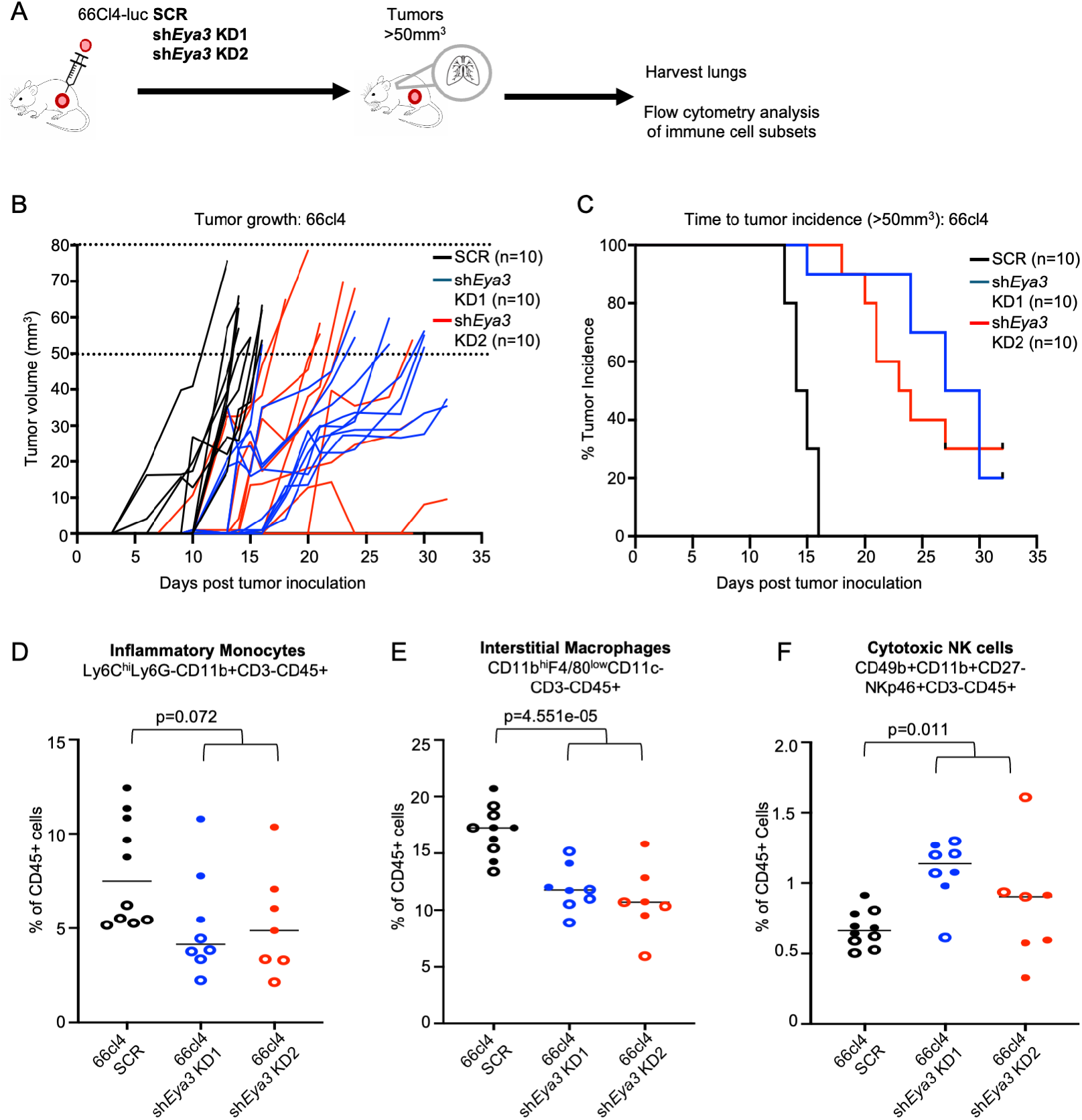
*Eya3* knockdown in triple negative breast cancer cells alters immune infiltration to the pre-metastatic lung. (A) Schematic representation of the experimental setup. 66cl4 SCR, sh*Eya3* KD1, and sh*Eya3* KD2 cells were orthotopically injected into the fourth mammary fat pad of syngeneic BALB/c mice. When tumor size exceeded 50mm^3^, mice were sacrificed. Lung single cell suspensions were analyzed for immune cells by flow cytometry. (B) Growth of pre-metastatic primary tumors as measured by caliper measurement. Mice that did not reach 50mm^3^ after ∼30 days were excluded from flow analyses. (C) Kaplan Meier curves of time for tumors to reach 50mm^3^. D-F). Flow cytometry analysis was used to examine the presence of inflammatory monocytes (D), interstitial macrophages (E), and cytotoxic NK cells (F) in the lungs of pre-metastatic mice. Data was collected from 7-10 mice per group, from 2 independent experiments (represented by open or closed circles). Mice that did not form tumors were removed from analysis. Statistical analysis was performed using ANOVA with sum contrasts in R.

Next, we analyzed the lung single cell suspensions from these mice by flow cytometry for the presence of various immune cell subsets (Suppl Fig. 6C). We did not observe significant changes in CD45+ lymphocytes overall, CD8+ T cells, CD4+ T cells, B cells, or neutrophils in the lungs of mice injected with SCR cells at the orthotopic site when compared to those injected with *Eya3* KD cells (Suppl. Fig. 6D). However, mice harboring small *Eya3* KD tumors demonstrated reduced levels of inflammatory monocytes (Fig. 4D) and interstitial macrophages (Fig. 4E) in the pre-metastatic lung compared to those with SCR tumors. In addition, we observed elevated levels of cytotoxic Natural Killer (NK) cells, marked by CD11b high/positive and CD27 low/negative^99^, in the lungs of mice injected orthotopically with *Eya3* KD cells when compared to SCR control cells (Fig. 4F). While we have previously observed changes to myeloid cells in the primary tumor with *Eya3* KD^25^, we did not observe any changes to cytotoxic NK cell levels in early primary tumors by flow cytometry (Suppl. Fig. 7A) or Vectra staining (Suppl. Fig. 7B). These data demonstrate that *Eya3* KD promotes the influx of anti-tumor cytotoxic NK cells to the pre-metastatic lung, while decreasing the infiltration of potential pro-tumor innate immune cells.

### EYA3 regulates secreted factors that inhibit cytotoxic NK cell infiltration and activation potential

Our data demonstrate that cancer cells in the primary tumor can influence immune cell residence in the pre-metastatic lung. Given that these effects are driven by the primary tumor in the absence of detectable micrometastases in the lung, we hypothesized that EYA3 must regulate secreted factors that mediate this effect. To test this hypothesis, we incubated phenol red and serum-free media with cancer cells for 48 hours, after which we injected this conditioned media (CM) into mice daily for 7 days (Fig 5A). After 7 days, we sacrificed the mice and analyzed immune cell infiltration into the lung. In the absence of a primary tumor, we did not observe significant differences in the infiltration of inflammatory monocytes or interstitial macrophages to the lungs of mice that received CM from *Eya3* KD cells compared to scramble cells (Fig. 5B, 5C). Overall, there were fewer inflammatory monocytes and interstitial macrophages in the lungs of tumor-free mice receiving CM compared to the lungs of mice bearing primary tumors (Fig. 5B, 5C, 4D, 4E), suggesting that the presence of the primary tumor is required for the changes we observed in these populations in our previous experiments.

**Figure 5:**
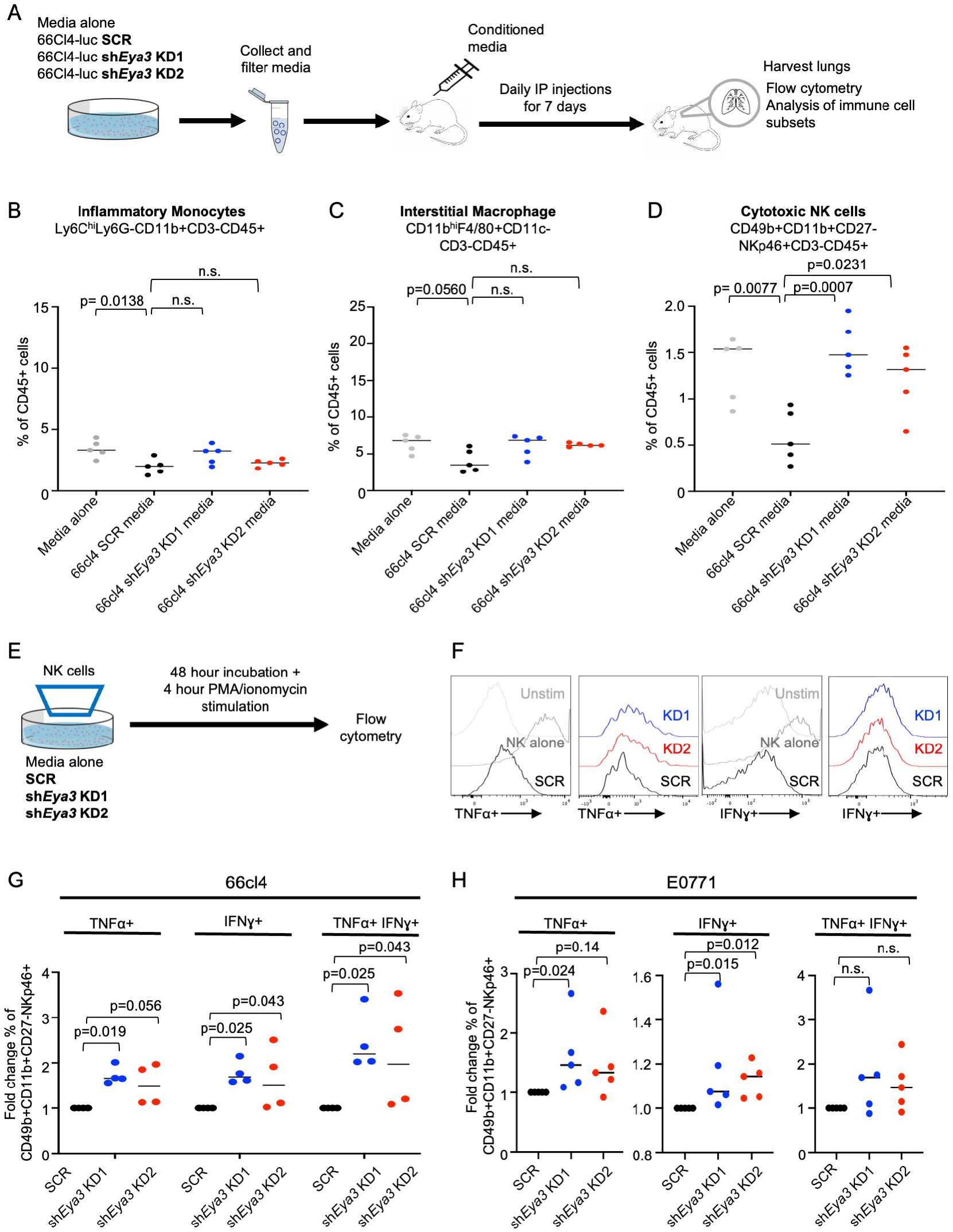
Cancer cell secreted factors alter cytotoxic NK cell infiltration to the lung and reduce cytotoxic NK cell activation *in vitro*. (A) Schematic representation of the experimental setup. Briefly, phenol red-free, serum-free media was incubated with cancer cells with or without *Eya3* KD in vitro for 48 hours, then collected and filtered. 300μL of media was injected intraperitoneally into mice for 7 days, then mice were sacrificed and lungs were made into single cell suspensions. (B-D) Lungs were analyzed for immune cell subsets by flow cytometry. Data were collected from 5 mice per group. (E) Schematic representation of the experimental setup. Briefly, NK cells were enriched from syngeneic mouse spleens using negative selection beads and incubated for 48 hours with IL-15 on a 0.4μM transwell placed on top of cancer cells with or without *Eya3* KD. PMA + ionomycin + golgi plug was added for the last 4 hours of incubation. (F-G) NK cell-enriched splenocytes cocultured with various 66cl4 cells were analyzed for the production of TNFα and IFNɣ by flow cytometry. Shown in F is a representative histogram of TNFα and IFNɣ flow cytometry staining. Gates were drawn based on unstained control, and % of cells expressing TNFα and IFNɣ were quantified and normalized to SCR. Data were collected from 4 independent experiments and graphed in G, and statistical analysis was performed using Kruskal-Wallis, adjusted for multiple comparisons. (H) As in G, for E0771. Data were collected from 5 independent experiments, and statistical analysis was performed using Kruskal-Wallis, adjusted for multiple comparisons.

However, we did observe a statistically significant increase in cytotoxic NK cells in the lungs of mice that received CM from 66cl4 *Eya3* KD cells when compared to mice that received media from 66cl4 SCR cells (Fig. 5D). We tested this result in a second cell line, E0771, and observed a similar trend towards increased cytotoxic NK cells in the lungs of mice that received CM from E0771 *Eya3* KD cells compared to E0771 SCR cells (Suppl. Fig 7E), without notable differences in inflammatory monocytes or interstitial macrophages (Suppl. Fig. 7C, 7D). Together, these results suggest that EYA3-regulated changes in inflammatory monocytes and macrophages require the presence of a primary tumor, while EYA3-regulated secreted factors alone influence the presence of cytotoxic NK cells in the lung, even in the absence of a primary tumor.

While these data demonstrate a role for EYA3 in altering cytotoxic NK cell numbers in the lung, they do not determine whether EYA3 can directly regulate NK cell function. To determine whether TNBC Eya3 can directly affect NK cell function, we enriched for NK cells from mouse splenocytes and cocultured them with BC cells with or without *Eya3* KD in a transwell system that allows for the exchange of secreted factors but does not allow for direct interaction between cancer cells and NK cells. We then stimulated the NK cell-enriched splenocytes with PMA and ionomycin to activate them and measured the ability of NK cells to produce the cytotoxic cytokines TNFα and IFNɣ by flow cytometry (Fig. 5E). Compared to NK cells cultured in the absence of TNBC cells, NK cells cocultured with 66cl4 SCR cells showed reduced ability to produce TNFα and IFNɣ (Fig. 5F). However, when NK cells were cultured with *Eya3* KD cells, we observed an increase in the percentage of activated cytotoxic NK cells producing TNFα and IFNɣ (Fig. 5F, 5G). We observed similar effects on cytokine production in a second breast cancer model, E0771, whereby a higher percentage of NK cells cocultured with E0771 *Eya3* KD cells were able to produce cytotoxic cytokines upon activation (Fig. 5H). These data demonstrate that secreted factors from EYA3-expressing cells reduce the ability of NK cells to produce cytotoxic cytokines upon activation, highlighting a role for EYA3 in directly regulating NK cell activation potential.

### EYA3 promotes expression of the cytokine CCL2 through activation of NF-kB signaling

Immune cell recruitment and activation are largely driven by cytokine signaling, and NF-kB signaling has a well-established role in regulating the expression of various cytokines. Thus, we hypothesized that Eya3-NF-kB-mediated cytokine expression is responsible for changes to the immune pre-metastatic niche. To test this hypothesis, we first reevaluated our RNA-seq data from *Eya3* KD cells with or without *IKK2-SE* addback (Fig. 3, Suppl. Fig. 3C) to interrogate changes in cytokine expression by using the GSEA GO Molecular Function Cytokine Activity dataset (Fig. 6A). We observed that only a subset of cytokines that were changed with *Eya3* KD were also rescued with *IKK2-SE* addback (Fig. 6A). We focused on cytokines that appeared to be positively regulated by EYA3-NF-kB activity (inset panel on Fig. 6A), and tested the effect of *Eya3* KD on the expression of these cytokines in multiple TNBC cell lines by qRT-PCR, including additional cytokines previously implicated in pre-metastatic niche formation in our analysis^62^ (Fig. 6B, Suppl. Fig. 8A, 8B). Using two independent *Eya3* shRNA constructs in 3 different mouse mammary carcinoma cell lines, we found that *Ccl2* was most consistently and statistically significantly changed with *Eya3* KD (Fig. 6B, 6C, Suppl. Fig. 8A, 8B). We also observed a reduction in secreted CCL2 protein with *Eya3* KD by performing Western blot analysis (Fig. 6D), and/or ELISA (Suppl. Fig. 8C, 8D) on cancer cell CM from several mouse TNBC lines.

**Figure 6:**
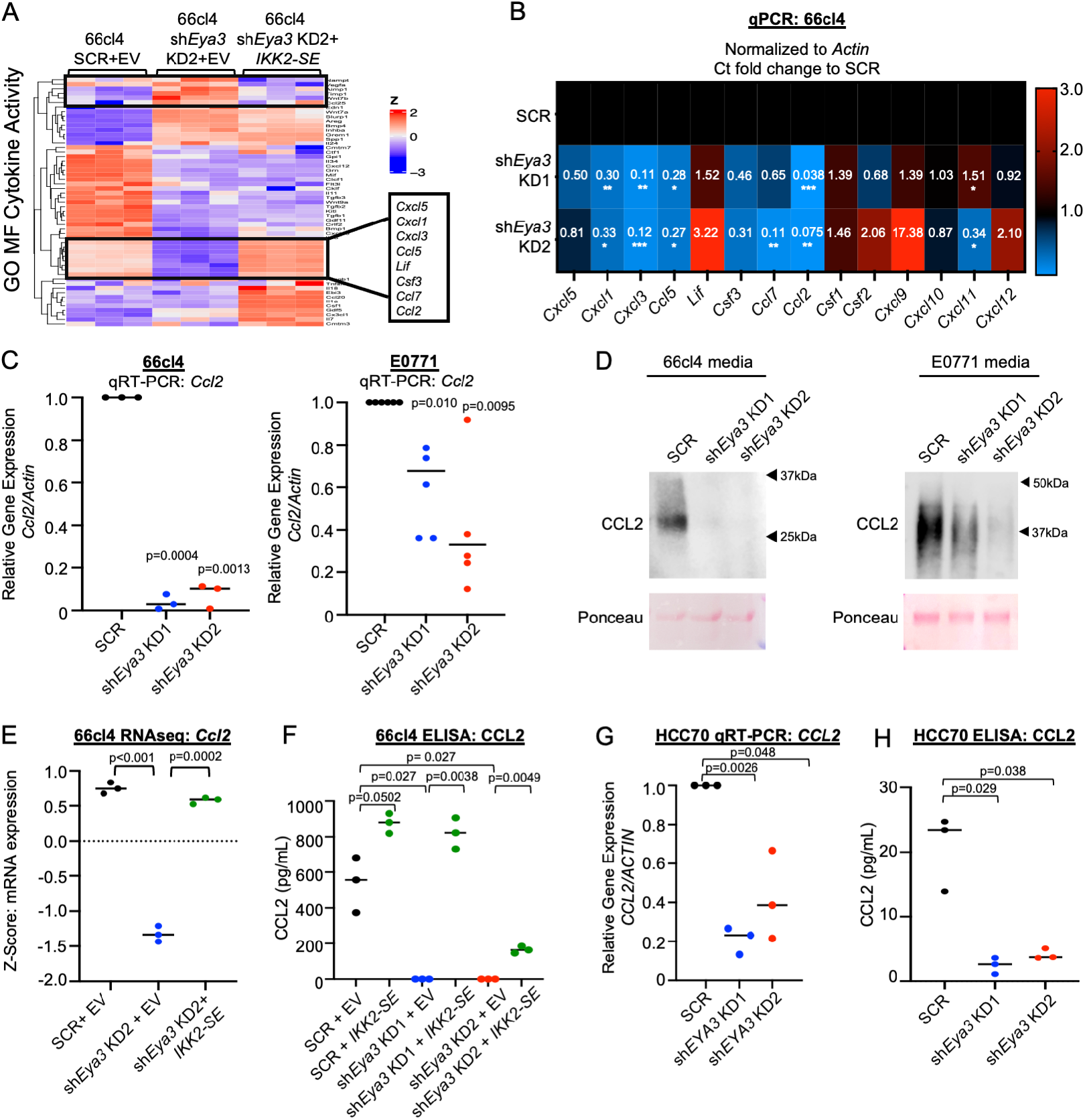
EYA3 promotes expression of the pre-metastatic niche regulator CCL2, downstream of NF-kB. (A) Heat map of DEGs in the GO MF (Molecular Function) Cytokine Activity gene set from RNA-sequencing of 66cl4 SCR+EV, sh*Eya3* KD2+EV and sh*Eya3* KD2+*IKK2-SE*. (B) 66cl4 cells with or without *Eya3* KD were probed for cytokine expression by qRT-PCR, normalized to *Actin*, and fold change was calculated compared to SCR cells. (C) 66cl4 or E0771 cells with or without *Eya3* KD were probed for *Ccl2* expression by qRT-PCR and normalized to *Actin*. Fold change was calculated compared to SCR cells. (D) Media was incubated with 66cl4 or E0771 cells for 48 hours. Conditioned media was then probed for CCL2 by Western blot. Shown is a representative image of three independent experiments. (E) Z-transformed mRNA expression for *Ccl2* was plotted between shSCR + EV, sh*Eya3* KD2 + EV, and sh*Eya3* KD2 + *IKK2-SE* from RNA-sequencing, related to (A). (F) Media was incubated with 66cl4 cells for 48 hours, after which CCL2 levels were determined by ELISA. (G) As in C, for HCC70 cells. (H) As in F, for HCC70 cells. Statistical analyses for all qRT-PCR and ELISA data were performed using one-way ANOVA assuming unequal variances; when the overall F test was found to be p<0.05, unpaired two-tailed Welch’s t-test was performed to assess differences between experimental predefined comparison groups. All experiments are representative of three independent experiments. *p<0.05, **p<0.01, ***p<0.001

In our RNA-seq dataset, *Ccl2* mRNA was dramatically decreased with *Eya3* KD, and rescued with *IKK2-SE* add-back (Fig. 6E), suggesting that *Ccl2* is regulated by EYA3 downstream of NF-kB activation. This finding was confirmed by examining CCL2 protein levels with *IKK2-SE* addback in *Eya3* KD cells by ELISA and Western blot (Fig. 6F, Suppl. Fig. 8E). In addition, treatment of 66cl4 cells with an IKK inhibitor, IKK inhibitor VII, led to a dose-dependent decrease in *Ccl2* mRNA and protein (Suppl. Fig. 8F, 8G), again demonstrating that CCL2 is downstream of EYA3/NF-kB. Importantly, we confirmed that the relationship between EYA3 and CCL2 could be found in human TNBC, as *EYA3* loss led to reduced CCL2 levels in human HCC70 cells (Fig. 6G, 6H; Suppl. Fig. 8H). These data identify CCL2 as a target of EYA3-NF-kB signaling in both mouse and human TNBC.

### *Ccl2* re-expression in *Eya3*-deficient cells rescues NK cell activation potential *in vitro* and cytotoxic NK cell infiltration to the lung

To determine whether CCL2 is responsible for EYA3 effects on cytotoxic NK cells, we utilized a retroviral approach to stably restore *Ccl2* expression to 66cl4 *Eya3*-KD cells. Cancer cell CM was collected from SCR control and *Eya3* KD cells +/- *Ccl2*, and re-expression of CCL2 protein was verified by WB (Fig. 7A) and ELISA (Fig. 7B). To test whether CCL2 is responsible for EYA3 effects on NK cell activation potential, we cocultured splenocytes enriched for NK cells with 66cl4 *Eya3* KD cells with or without *Ccl2* re-expression, stimulated cells with PMA and ionomycin, and tested for NK cell production of cytotoxic cytokines by flow cytometry (as outlined in Fig. 5E). We observed that *Ccl2* re-expression reduced the percentage of cytotoxic NK cells producing TNFα, IFNɣ, or both cytokines (Fig. 7C). Next, we tested the effects of CCL2 on NK cell infiltration to the lung. To this end, CM from each condition was injected into mice daily for 7 days (as outlined in Fig. 5A), after which mice were sacrificed and immune cell infiltration into the lungs was analyzed. Consistent with previous experiments (Fig. 5D), we observed an increase in cytotoxic NK cells in the lungs of mice that received CM from *Eya3* KD cells when compared to SCR cells (Fig. 7D), and these effects were partially rescued with *Ccl2* re-expression (Fig. 7D). Interestingly, *Ccl2* overexpression in SCR cells had no effect on cytotoxic NK cell residence in the lungs. Similar to previous experiments (Fig. 5B, 5C), we did not observe notable changes in interstitial macrophages or inflammatory monocytes (Suppl. Fig. 9A, 9B). These results demonstrate that CCL2 partially contributes to the effects of cancer cell-expressed EYA3 on NK cell cytokine production *in vitro* and NK cell infiltration in the lungs *in vivo*.

**Figure 7:**
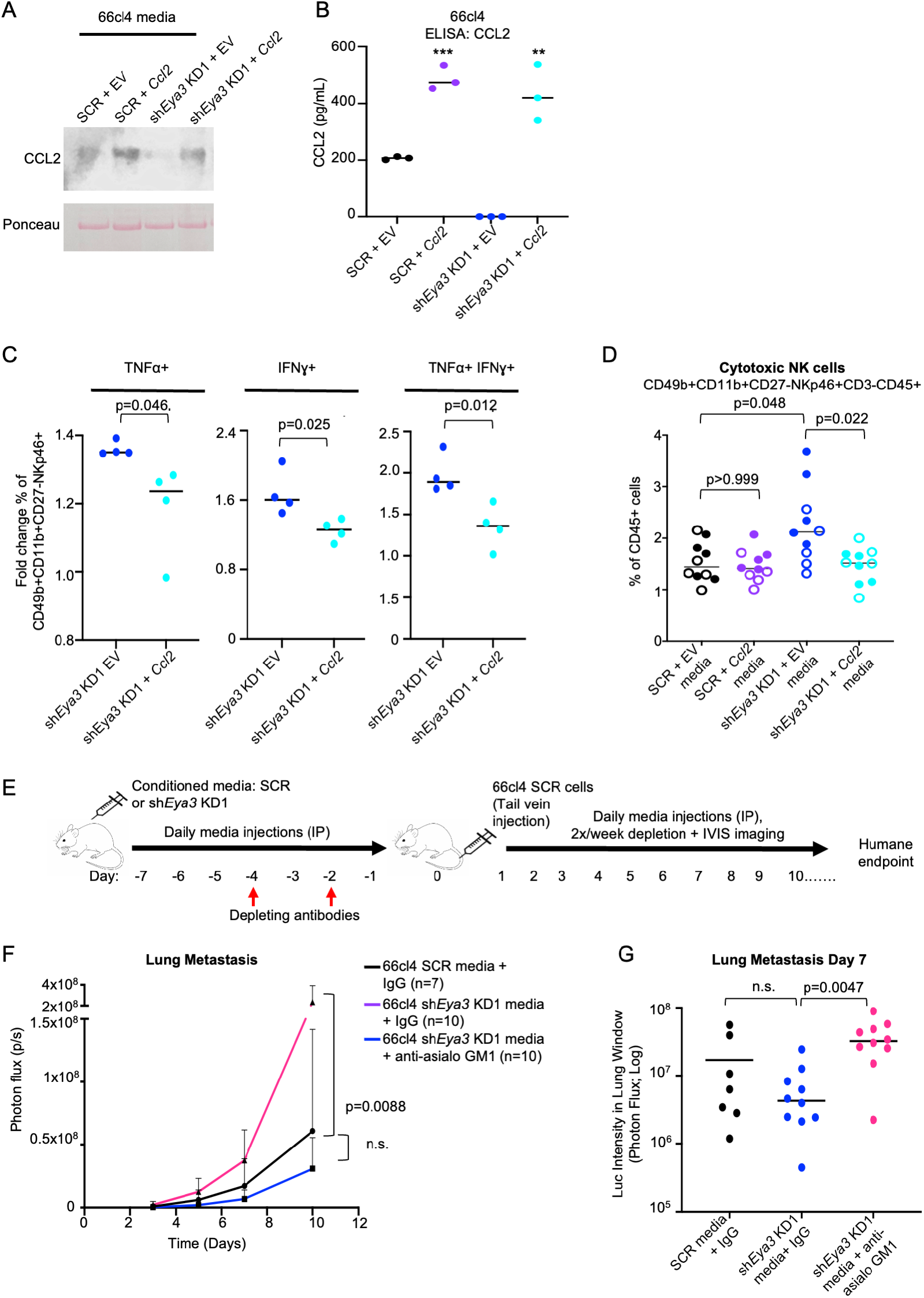
Expression of *Ccl2* in *Eya3* KD cells rescues effects of CM on cytotoxic NK cell infiltration to the lung and activation *in vitro*. (A-B) Retrovirus was used to stably express *Ccl2*, and media was analyzed for CCL2 levels by (A) Western blot and (B) ELISA assay. (C) NK cell-enriched splenocytes cocultured with 66cl4 cells were analyzed for production of TNFα and IFNɣ by flow cytometry. Data were collected from 4 independent experiments, and statistical analysis was performed using unpaired two-tailed Welch’s t-test. (D) Cytotoxic NK cells were analyzed by flow cytometry in the lungs of mice treated with CM. Data were collected from 10 mice per group, from two independent experiments (represented by open or closed circles). Statistical analysis was performed using Kruskal-Wallis, adjusted for multiple comparisons. (E) Schematic of the experimental setup. Briefly, CM was collected from cancer cells with or without *Eya3* KD after 48 hours of incubation. 300μL of CM was injected intraperitoneally into mice for 7 days prior to the injection of 66cl4 SCR cells via tail vein into all groups. Anti-asialo GM1 depleting antibody or IgG control was administered 4 and 2 days prior to cancer cell injection. Depletion was verified one day prior to cancer cell injection by flow cytometry analysis of blood. Mice were imaged 3x/week, depleting antibodies were administered 2x/week, and CM was injected daily throughout the course of the experiment. (F-G) Bioluminescence intensity in the lung window was detected by IVIS imaging and plotted over time. 3 mice were excluded from the SCR + IgG group due to poor tumor injection (marked by tail vein burst or complete absence of metastatic cells throughout the course of the experiment) or immediate death upon tumor injection. Statistical analysis for F was performed using longitudinal mixed model analysis in R, and for H using Kruskal-Wallis, adjusted for multiple comparisons.

To determine whether EYA3/NF-kB/CCL2 regulation of NK cells alters metastasis, we then injected cancer cell-CM from SCR control or *Eya3* KD cells into mice daily for 7 days to prime the pre-metastatic lung (Fig. 7E). During this time, NK cells were depleted from mice by administering anti-asialo GM1 antibody (used because BALB/c mice do not express the NK1.1 antigen^100^) or an IgG control, and depletion was verified by flow cytometry performed on blood drawn from treated mice (Suppl. Fig. 9C). 66cl4 SCR cells were then injected via tail vein into mice from all groups: SCR CM + IgG; (2) sh*Eya3* CM + IgG; (3) sh*Eya3* CM + anti-asialo GM1 (Fig. 7E). In this experiment, a modest decrease in metastatic colonization was observed in mice that were primed with sh*Eya3* CM compared to those that received SCR CM (Fig. 7F, 7G). Given that macrometastases were detectable in the lungs after only 3 days, and that 66cl4 SCR cells were injected (which are capable of producing their own CCL2), it is likely that this experimental setup gives low resolution for observing differences between the conditions treated with *Eya3* KD vs SCR CM. Nonetheless, depletion of NK cells led to a significant increase in lung metastatic colonization, and overall survival was greatly reduced (Fig. 7F, 7G, Suppl. Fig. 9D). These data demonstrate that NK cells play a critical role in preventing TNBC colonization in the lung, downstream of EYA3.

### EYA3 is associated with NF-kB activity and CCL2 expression in human breast tumors

To determine if our findings in mouse models are conserved in human breast cancer, we performed pairwise correlation analysis between the expression of *EYA3* and the expression of every other sequenced gene across over 2,500 patient tumors from the METABRIC dataset of invasive breast carcinomas (Fig. 8A). From this pairwise correlation analysis, we were able to generate a ranked list of genes with the strongest correlation with *EYA3* expression across all patients in this dataset. Using this list, we performed GSEA analysis to identify gene sets exhibiting a strong correlation with *EYA3* in these patient tumors. In support of our results in the mouse BC models, *EYA3* expression in these tumors was strongly associated with Hallmark TNFα Signaling via NF-kB and Hallmark Inflammatory Response gene sets (Fig. 8A; Supp. Fig. 10A, 10B), as well as with MYC target gene sets, aligning with the known function of EYA family members as regulators of MYC expression and stability^20,21^. In the same dataset, we observed a positive correlation between *CCL2* and *EYA3* mRNA levels in human breast cancer cell patient samples (Fig. 8B). For each patient sample, we calculated a score for the Hallmark

**Figure 8:**
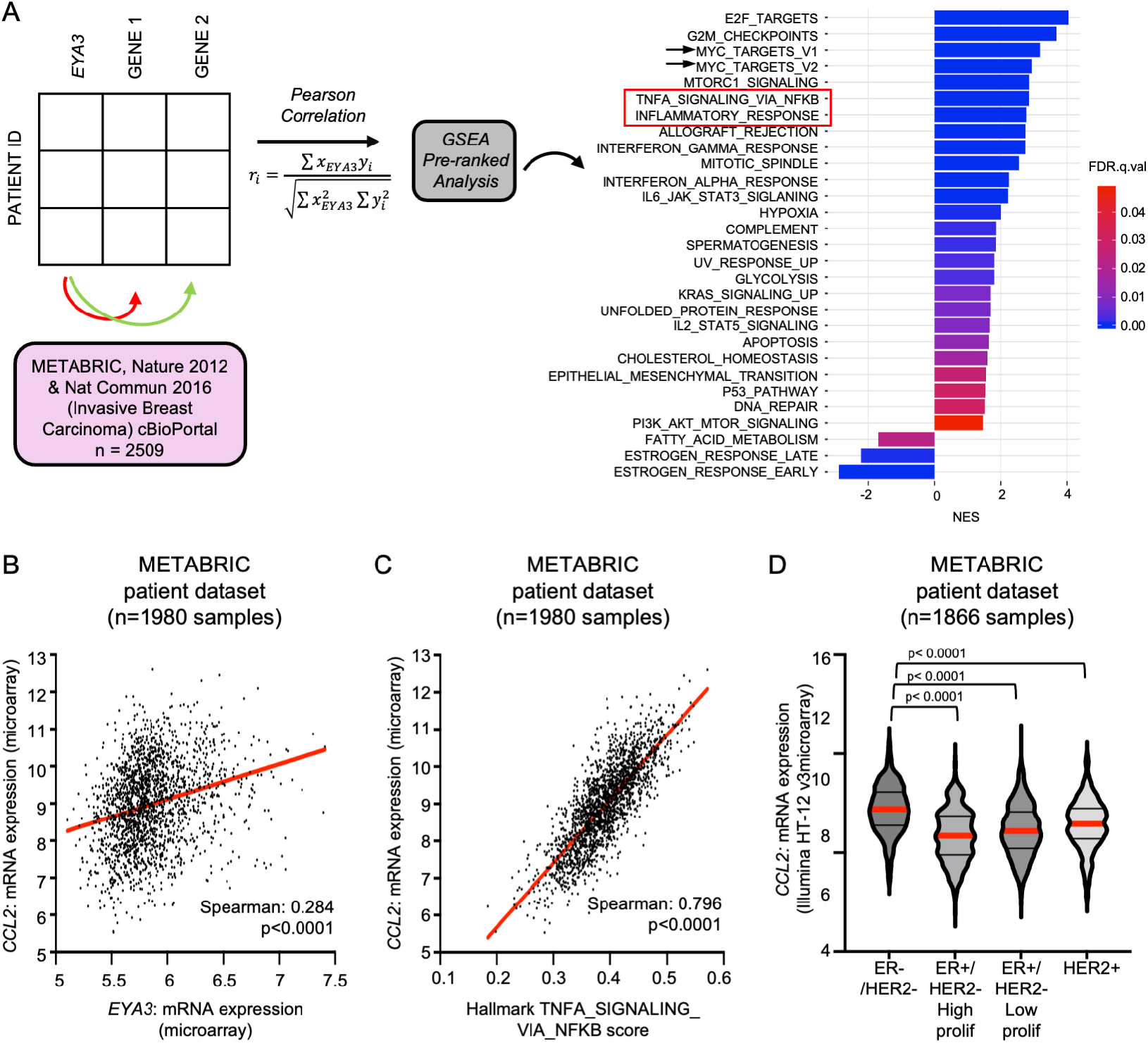
*EYA3* expression correlates with NF-kB signaling and *CCL2* expression in human breast cancer. (A) Schematic of pairwise correlation analysis of METABRIC patient data^80–82^ ranking correlated genes with *EYA3* and looking at enrichment of correlated genes within Hallmark gene sets. (B) Gene expression for the METABRIC dataset^80–82^ was downloaded from cBioPortal, and *CCL2* mRNA expression was compared against *EYA3* mRNA expression for each sample. (C) Using the GSVA R package, ssGSEA was used to calculate the NFkB score using the Hallmarks gene set collection, HALLMARK_TNFA_SIGNALING_VIA_NFKB^116,117^ for each sample in the METABRIC dataset and compared to *CCL2* mRNA expression. (D) *CCL2* mRNA expression was evaluated in varying genetic subsets in the METABRIC dataset.

TNFα Signaling via NF-kB signaling pathway and also observed a positive correlation between this pathway and *CCL2* levels (Fig. 8C). Interestingly, *CCL2* levels were found to be highest in the ER-/HER2- gene classifier subtype, suggesting that EYA3-NF-kB regulation of CCL2 expression may be most relevant to TNBC. We additionally observed a positive correlation between EYA3 and CCL2 mRNA in a second patient dataset from the Metastatic Breast Cancer project (Suppl. Fig. 10C). Together, these data demonstrate that EYA3, NF-kB pathway, and CCL2 are significantly correlated in samples from patients with breast cancer, supporting the existence of an EYA3- NF-kB-CCL2 signaling axis in human patients.

## Discussion

Understanding the mechanisms that drive metastatic progression is critical to developing new clinical strategies for patients with metastatic TNBC. Given that embryonic programs are coopted by cancer cells during tumor progression, we interrogated the role of the embryonic protein EYA3 in TNBC metastasis, and show for the first time that loss of EYA3 reduces the ability of TNBC cells to complete the full metastatic cascade from a primary tumor. We discover that EYA3 promotes NF-kB activation and subsequent CCL2 expression in multiple models of TNBC, and that EYA3/NF-kB/CCL2 signaling suppresses cytotoxic NK cell responses in the PMN, which are critical for metastasis. Altogether, this study elucidates tumor cell autonomous and non-cell autonomous mechanisms by which EYA3 promotes TNBC metastasis.

EYA proteins and NF-kB both drive key processes during embryonic development^12,101^, and our data demonstrate that these two developmental pathways intersect in cancer to promote TNBC metastasis. We show that reactivation of NF-kB signaling is sufficient to restore metastasis in *Eya3* KD cells, and also observe an increase in migration, which is in line with previous reports that tumor intrinsic NF-kB signaling promotes invasive phenotypes^92^. EYA3 enhances primary tumor growth by regulating a MYC/PDL1 axis^25^, but, intriguingly, we find that neither MYC signaling nor primary tumor growth is restored with NF-kB reactivation. Thus, in this study we discover that EYA3 differentially regulates MYC and NF-kB signaling to promote different stages of tumor progression. In mouse TNBC cells, we observe that EYA3 is predominantly in the cytoplasm (Fig. 2), suggesting that EYA3 may be regulating NF-kB signaling through its phosphatase activities rather than its transcriptional activities. In support of this idea, it has been demonstrated that in response to viral stimulation, EYA4 enhances innate immune signaling via STING and/or NF-kB in mouse embryonic fibroblasts through its threonine phosphatase activity^37^. EYA3 regulates MYC stability through association with the serine/threonine phosphatase PP2A-B55α^21^, and future studies should evaluate whether this same phosphatase activity drives NF-kB regulation and metastasis, or whether EYA3 regulates these two pathways through completely independent functions.

Cancer cells remodel the PMN to allow for more efficient colonization and metastatic outgrowth at secondary sites. From the primary tumor, cancer cell secreted factors alter the immune microenvironment in the PMN^98^ by influencing the recruitment, differentiation, and activity of various immune cells including monocytes^59^, macrophages^102^, neutrophils^58,64^, and NK cells^58^. While all of these cell types contribute to immunosurveillance at metastatic sites, NK cells are critical for eliminating metastatic cancer cells^65,98^, and cancer cell resistance to NK cell responses enables metastatic colonization and outgrowth^66^. CCL2 has been found to promote monocyte migration and macrophage retention in the PMN, resulting in enhanced metastasis^59,102^. However, the effect of CCL2 on NK cells in the PMN remains poorly understood. In this study, we uncover a previously unappreciated role for CCL2 in *negatively* regulating cytotoxic NK cells in the pre-metastatic lung. In the context of pregnancy, stromal-derived CCL2 inhibits cytotoxic NK cell cytotoxicity through activation of SOCS3^63^. On the contrary, in other settings CCL2 has been shown to directly *promote* NK cell migration and cytolysis. Given that NK cells adopt different phenotypes in different organs depending on local cues and interactions, it is not surprising that CCL2 may have disparate effects on NK cells in distinct contexts. Thus, the specific effect of CCL2 on cytotoxic NK cells in the pre-metastatic lung warrants further investigation.

Increasingly, NK cells have been highlighted for their potential in cancer immunotherapy, with large-scale efforts to develop therapies that boost NK cell responses or create “off-the-shelf” NK cells for adoptive cell therapy^68^. Emerging evidence also suggests that NK cells positively contribute to the response to PD-1/PD-L1 inhibitors^103^, which are already FDA-approved for use in metastatic TNBC. Understanding how tumor intrinsic mechanisms inhibit NK cell-mediated immunosurveillance is critical to the optimization of these therapeutic strategies.

Due to its myriad functions in cancer cells, NF-kB is an attractive therapeutic target. However, the ubiquity of this signaling pathway and various *anti*-tumor activities of NF-kB make systemic therapeutic approaches challenging. In the same vein, CCL2 may be difficult to target given its diverse roles in immune cell signaling. In this study we demonstrate that EYA3 is upstream of NF-kB/CCL2, presenting EYA3 as a promising target for inhibition of this pathway. EYA3 is highly expressed in TNBC, suggesting that EYA3 inhibition could more specifically reduce NF- kB activation in TNBC cells, likely with low side-effect profiles given that knockout of EYA3 in mice does not induce any major phenotypes^104^. EYA inhibitors are already in existence^105^, supporting the notion that EYA proteins can be pharmacologically targeted. However, many of these inhibitors target EYA phosphotyrosine activity, and new inhibitors may need to be developed if a different EYA3 phosphatase activity (such as that mediated through its interaction with PP2A^21^) is implicated in NF-kB/CCL2 regulation.

In this study we uncover the existence of a previously unappreciated EYA3/NF-kB/CCL2 signaling axis in TNBC metastasis, and demonstrate that this pathway suppresses cytotoxic NK cells in the PMN to promote metastasis. Moreover, our past findings that EYA3 promotes MYC stabilization and suppresses CD8+ T cells in the primary tumor demonstrate that EYA3 plays numerous roles in TNBC progression. Our data suggest that EYA3 inhibition will impact not only primary tumor growth but also reduce metastasis, and positively impact both adaptive and innate immune responses at both sites of cancer cell growth. Taken together, these data suggest that targeting EYA3, particularly in combination with immunotherapies, may be a powerful means to inhibit TNBC progresson.

## Acknowledgments

This work has been supported through grants from the NIH: R01 R01CA224867, R01CA221282, R01CA275187 (HLF), K00CA245552 (to SRR) and training fellowships F30CA257215 (to CJH) and F31CA275314 (to AW). Special thanks to Andrew Goodspeed, PhD and Etienne Danis, PhD for RNA-sequencing data processing. Thank you to Patricia Ernst, PhD for subcloning and providing aliquots of the pLVX-EF1α-IRES-mCherry empty vector and IKK2-SE plasmids. We thank the Human Immune Monitoring Shared Resource (RRID:SCR_021985) within the University of Colorado Human Immunology and Immunotherapy Initiative and the University of Colorado Cancer Center (P30CA046934) for their expert assistance in analysis of Vectra staining. This work also utilized the Cell Technologies (RRID:SCR_021982), Genomics (RRID:SCR_021984), Biostatistics and Bioinformatics (RRID:SCR_021982), and Flow Cytometry (RRID:SCR_022035) shared resources at the University of Colorado Anschutz Medical Campus and the University of Colorado Cancer Center (P30CA046934).

## Methods

### METABRIC survival analysis

Gene expression and survival data for the METABRIC dataset^80–82^ were downloaded from cBioPortal (May 30, 2024). Gene expression for EYA3 was used to stratify the overall cohort into quartiles. Individuals from the top and bottom quartiles were compared for overall survival using the survival (v3.5-8) and survminer (v0.4.9) R packages. Overall survival was analyzed with patients living past 10 years right censored; the log-rank test was used to calculate statistical significance.

### Cell lines and culture conditions

TNBC cell line 66cl4 was obtained as a generous gift from Fred Miller^106^, and cultured in DMEM High Glucose media supplemented with 10% calf serum, 1% L-glutamine, 1% Pen/Strep and 1% non-essential amino acids. The Met1 murine mammary carcinoma cell line was obtained as a generous gift from Jeffrey Greg^107^, cultured in DMEM High Glucose media supplemented with 10% fetal bovine serum (FBS), 1% L-glutamine,1% Pen/Strep. E0771 gfp-luc murine breast cancer cells were cultured in RPMI supplemented with 10% FBS and 1% Pen/Strep. The human TNBC cell line HCC70 (purchased from the Cell Technologies Shared Resource on campus, STR profiled in April 2010 immediately prior to freezing down, and obtained originally from ATCC) was cultured in DMEM + F12 Ham’s media supplemented with 10% FBS + 0.5% Pen/Strep. All cell lines were grown at 37°C in 5% CO2. All cell lines are checked for mycoplasma every 3-6 months to ensure accuracy of our findings. Cell lines were passaged at least 2 times after thawing prior to use in experiments, and were re-thawed periodically to maintain low passage numbers.

### Generation of knockdown cell lines

Eya3 was knocked down in murine cell lines using two different shRNAs (shEya3 KD1 - TRCN0000029858 and shEya3 KD2 - TRCN0000029855, Dharmacon, Lafayette, CO) and control scramble shRNA (Addgene plasmid 1864, Cambridge, MA). ShRNAs were lentivirally introduced into cells according to the PLKO.1 manufacturing protocol (Addgene, Cambridge, MA). All cells were maintained in 2.5 ug/ml puromycin for selection of pooled populations and maintenance of knockdown. Parental cells were tagged with firefly luciferase using the MSCV Luciferase PGK-hygro vector (Addgene plasmid #18782, Cambridge, MA) and transduced cells were selected as a pooled population with hygromycin (200µg/mL). E0771 cells were previously tagged with firefly luciferase using MSCV Luciferase PGK-hygro vector (Addgene plasmid #18782) and shEya3 KD cells were generated using the above Eya3 KD shRNAs.

### Animals and *in vivo* studies

All animal studies were performed according to protocols reviewed and approved by the Institutional Animal Care and Use Committee (IACUC) at the University of Colorado AMC. Only female mice were used in this study because breast cancer predominantly affects females (>99%). Animals were randomly assigned to experimental groups.

For spontaneous metastasis studies, 200,000 cells in Opti-MEM Reduced Serum Media (Gibco A4124801) were injected into the fourth mammary fat pad of 8-10 week old syngeneic BALB/cJ mice (Jackson Laboratories RRID:IMSR_JAX:000651) or three- to twelve-week-old male NOG/SCIDγ (NSG) mice (bred in-house). For IKK2-SE addback studies, 250,000 cells were used as these additionally carry an mCherry tag. Tumors were measured with calipers at indicated intervals and volume was calculated as [long axis (cm)*short axis^2^ (cm)]/2. Once a tumor surpassed a volume of 1cm^3^ (unless otherwise indicated), primary tumors were surgically removed. Mice that died secondary to surgical complications were excluded from survival analyses. One week after surgery, mice were imaged weekly under anesthesia on the IVIS200 bioluminescent imaging system 10 minutes after receiving an intraperitoneal injection of 100µl of 100X luciferin (Gold Biotechnology, LUCK-1G). All mice were sacrificed upon clinical signs of health deterioration, confirmed by a veterinarian. Bioluminescent images were analyzed using Living Image software. All images were normalized based on photon flux, and lung luciferase intensity was determined through selection of a lung window with Living Image software to capture luciferase signal around the lung.

For tail vein injection studies, 500,000 luciferase-gfp tagged E0771 cells were injected intravenously into 8-10 week old syngeneic C57BL/6J mice (Jackson Laboratories). After 3 days, mice were imaged every 3-4 days with the IVIS200 bioluminescent imaging system as described above. Mice were monitored for labored breathing and weight loss, and began reaching humane endpoint for sacrifice on day 21; all mice were sacrificed by day 23. Lungs were collected into RNAlater RNA preservation solution (Sigma-Aldrich), and frozen at −80. Lung tissue was subsequently minced and mixed, and RNA was isolated for analysis of Puro and Actin expression by qRT-PCR.

For pre-metastatic niche studies, 200,000 cells were injected into the fourth mammary fat pad and tumor growth was tracked by caliper measurement. Tumors were considered palpable when they exceeded 50mm^3^, at which point mice were sacrified for downstream analyses. For media injection studies, cancer cells were plated in 6 well plates and the following day media was changed to 2mL/well of phenol red-free, serum-free high glucose DMEM supplemented with L-glut and non-essential amino acids. After 48 hours, media was collected, passed through a 0.2μm filter, and frozen at −80°C. Mice were injected intraperitoneally with 300μL conditioned media each day for 7 days, then sacrified and lungs were collected.

For NK cell depletion studies, animals were treated with 300μL conditioned media daily for 7 days prior to injection of 200,000 66cl4 cells into the tail vein. Conditioned media was injected daily throughout the duration of the experiment. For depletion of NK cells, 50μg of anti-asialo GM1 antibody (Invitrogen, #16-6507-39) or 50μg polyclonal Rabbit IgG (BioXcell, #BE0095) was injected intraperitoneally 4 days prior to tumor cell injection, 100μg injected 2 days prior to tumor cell injection, and then 50μg injected twice weekly for the duration of the experiment. Mice were imaged 3 times per week by IVIS200 bioluminescent imaging as above.

### RNA expression analysis

Total RNA was extracted from cells at 80% confluency using the RNeasy Plus Mini RNA Isolation Kit (Qiagen, 74136) according to manufacturer’s instructions. cDNA synthesis was performed using iScript cDNA Synthesis Kit (Biorad, 1708891) from 1µg of mRNA. qRT–PCR assays were performed using ssoFast Evagreen Supermix (BioRad, 1725205) and analyzed using the Biorad CFX96 qPCR instrument. The primer sequences used are listed in the supplemental information.

### Western blot analysis

Whole cell protein lysates (WCL) were generated by lysis in radioimmunopreciptation (RIPA) buffer supplemented with protease inhibitors and lysed via sonication. For Western blot analysis of cancer cell media, serum-free conditioned media was mixed at a 1:1 ratio with 2x Laemmeli buffer. Cytoplasmic/Nuclear fractionated protein extracts were generated via the NE-PER Nuclear and Cytoplasmic Extraction kit (Thermo Scientific, 78835) according to the manufacturer’s instructions. Protein quantification was determined via Lowry assay. For WCL and Cytoplasmic/Nuclear protein immunoblots, 25-50μg of protein were electrophoresed on 5% stacking and 10% running gels or gradient gels and transferred to PVDF membranes.

Membranes were incubated for 1 hour with Ponceau stain and then imaged prior to blocking, followed by incubation overnight in primary antibody at 4°C then 1 hour at RT in secondary antibody prior to imaging. Blots were developed using SuperSignal West Pico Chemiluminescent Substrate (Thermo Scientific, 34080) and/or SuperSignal West Femto Chemiluminscent Substrate (Thermo Scientific, 34096) and imaged on the Licor Odyssey FC. For re-imaging of different proteins at similar molecular weights, blots were stripped with Restore Western Blot Stripping Buffer (Thermo Scientific, 21059) for 5 minutes at RT, then were blocked and re-probed as described above. Primary antibodies (all diluted in 5% BSA in TBST) are listed in supplemental information.

### RNA-Sequencing Analysis

RNA was extracted from samples using the RNeasy RNA Isolation Kit (Qiagen) in triplicate and sent to the University of Colorado Cancer Center Genomics Core Facility for sample preparation, sequencing and generation of raw data. Library preparation was performed using the NuGEN Universal Plus mRNA-Seq kit. Samples were sequenced with paired-end reads (40 million reads/sample) on the Illumina NovaSEQ6000 sequencing platform. The quality of the fastq files was assessed using FastQC^108^ and MultiQC^109^. Illumina adapters and low-quality reads were filtered out using BBDuk (http://jgi.doe.gov/data-and-tools/bb-tools). Trimmed fastq files were aligned to the hg38 human reference genome and aligned counts per gene were quantified using STAR^110^. Transcript expression levels were estimated with Salmon using inferential replicates^111^. Differential gene and transcript analyses were performed using the DESeq2^112^ and Swish^113^ packages. After processing, the University of Colorado Biostatistics and Bioinformatics Shared Resource RNA-seq analysis tool was used to perform pathway analysis, including over-representation analysis, functional gene set enrichment analysis (GSEA), transcription factor motif enrichment analysis, and Kegg Topology analysis. A fold change cutoff of 2 was used for transcription factor motif enrichment analysis.

### Generation of 66cl4 NF-kB and *Ccl2* rescue cell lines

Previously generated 66cl4 SCR were transduced with a pLVX-EF1a-IRES-mCherry empty vector (EV) construct and 66cl4 shEya3 KD2 cells were transduced with either the EV or a pLVX-EF1a-IRES-mCherry IKK2-SE construct (both constructs generously donated by Patricia Ernst, PhD). After transduction, cells were sorted using flow cytometry for mCherry positivity. mCherry-positive cells were used for all subsequent experiments – cells were re-sorted every 2- 4 weeks in culture to ensure high mCherry levels. For Ccl2 expression, mouse Ccl2 was amplified from mouse cDNA and cloned into pMSCVneo (Clonetech #631461, Mountain View, CA). The pMSCVneo-Ccl2 construct and retroviral packaging plasmid pcL-Eco were co-transfected into HEK293T cells using FuGENE transfection reagent (Promega, Madison, WI) to generate retroviral particles. Cells were transduced with viral particles and selected with 500μg/mL neomycin.

### Flow cytometry

After mice were humanely sacrificed, the lungs were perfused with 10mL of PBS and placed immediately into 1.5mL immune cell media (RPMI + 10% FBS + 1% Pen/Strep + 50μM β-mercaptoethanol) on ice. Lungs were then removed from media, minced, and incubated in digestion buffer (1x HBSS, 0.1mg/ml Collagenase IA, 60 U/ml DNase I) at 37°C for 25 minutes with continuous shaking. Cells were washed with medium and filtered through a 70μm nylon filter, then frozen in complete medium with 10% DMSO at -80°C. Lung single cell suspensions were thawed and resuspended in immune cell media and 1/10 of each lung was distributed into a round-bottom 96 well for each flow cytometry panel. For analysis of lymphocytes in the blood, 100μL-200μL of blood was collected by submandibular bleed into 500μL 2μM EDTA-HBSS, and red blood cells were lysed with 1X ACK buffer for 5mins prior to subsequent staining.

Cells were incubated with Zombie Fixable Viability Dye (BioLegend #423101, San Diego, CA) for 10 mins, washed and then stained for 30mins with fluorochrome-conjugated antibodies. Cells were then fixed using BD Cytofix/Cytoperm™ Fixation/Permeabilization Kit (BD Biosciences #554714, Franklin Lakes, NJ) or fixed and nuclear permeabilized using the eBioscience™ FOXP3/Transcription factor buffer staining set (eBioscience #00-5523-00, San Diego, CA). For analysis of T regulatory cells, cells were then incubated with an antibody specific for Foxp3 after permebaolization. For analysis of cytokine production, cells were incubated with antibodies specific for IFNƔ and TNFɑ after permebolization. Stained cells were analyzed on a Biorad ZE5 Cell Analyzer in the University of Colorado Cancer Center Flow Cytometry Shared Resource using Flowjo software (TreeStar, Ashaland, OR). Antibodies used for these analyses are listed in supplemental information.

### NK cell coculture assay

Spleens from C57BL/6 or BALB/c mice were passed through a 70μm nylon filter to make a single cell suspension. EasySep™ Mouse NK Cell Isolation Kit (STEMCELL Technologies #19855, Vancouver, Canada) was used to isolate mouse NK cells from spleen single cell suspensions according to the manufacturer’s instructions, and NK cell enrichment was verified by flow cytometry staining. Cancer cells were plated into a 12-well plate, and the following day the media was changed to serum-free cancer cell media. NK cells were then seeded in immune cell media (supplemented with fresh 10ng/mL mouse IL-15) onto a 12mm Transwell with 0.4μm Pore Polyester Membrane insert placed over top of cancer cells. Cells were co-incubated for 48 hours, and in the last 4 hours, 50ng/mL phorbol myristate acetate (PMA), 1μg/mL ionomycin, and 1X (3μg/mL) Brefeldin A golgi plugs were added to NK cells. NK cells were then transferred to a round-bottom 96-well plate and stained for flow cytometry.

### ELISA assays

Cancer cells were incubated with serum-free media for 48 hours. Media was collected and centrifuged to remove any floating cells. ELISA was performed on conditioned media using Mouse or Human MCP1 ELISA kit (Abcam #ab100721 and #ab100586, Cambridge, United Kingdom) according to the manufacturer’s instructions.

### Correlation analyses from patient datasets

For pairwise-correlation GSEA analysis, we accessed patient RNA expression data from the METABRIC patient cohort^80,81^ via access through cBioPortal^114,115^ as z-scores of gene expression and performed a pairwise Pearson correlation between EYA3 and all other genes in the dataset using R (4.0.1) and R Studio (v.1.2.5033) and subsequently ranked genes based on their correlation with EYA3 expression. We used the ranked list as input for pre-ranked Gene Set Enrichment Analysis (v4.0.3) using the Hallmarks Molecular Signature Database (h.all.v7.1) and identified pathways enriched with genes correlated with EYA3 expression in patient breast tumors. We plotted the significantly up- and down-regulated pathways using the ggplot2 package.

For EYA3 mRNA correlations, gene expression data for the METABRIC dataset^80–82^ were downloaded from cBioPortal (May 30, 2024). Using the GSVA (v1.48.3) R package, ssGSEA was used to calculate the NFkB score using the Hallmarks gene set collection (v2023.2), HALLMARK_TNFA_SIGNALING_VIA_NFKB^116,117^. Patients were stratified by 3- Gene classifier subtype^80,81^ on cBioPortal. Data was plotted in graphpad prism, and Spearman correlation was calculated for each correlation. EYA3 and CCL2 mRNA data from The Metastatic Breast Cancer Project were anayzed on cBioPortal.

## Statistics

Prism software (v9.0; GraphPad) was used for all statistical analyses with the exception of longitudinal mixed-model analyses described below. In all *in vitro* experiments, the conditions were run with at least 3 replicates and repeated at least 3 times. Two-tailed unpaired Student’s *T*-test was used when only a pair of conditions was compared. One-way or two-way Analysis of Variance (ANOVA) parametric followed by Bonferroni or Dunnett’s post-hoc test was used for multiple comparisons between control and knockdown conditions. Experiments demonstrating changes over time (cell growth, tumor growth, luciferase intensity curves and metastasis assays) were compared using a longitudinal mixed effects model that compares groups over repeated measures. All survival curves were compared using log-rank (Mantel-Cox) analysis.

## Data Availability Statement

The data supporting the findings of this study are available within the paper and its Supplementary Information. Raw NGS data will be deposited into the Gene Expression Omnibus and will be publicly available at the time of publication (GEO accession numbers will be provided at this time).

