## Supplemental information for "An EYA3/NF-κB/CCL2 signaling axis suppresses cytotoxic NK cells in the pre-metastatic niche to promote triple negative breast cancer metastasis"

Suppl. Figure 1

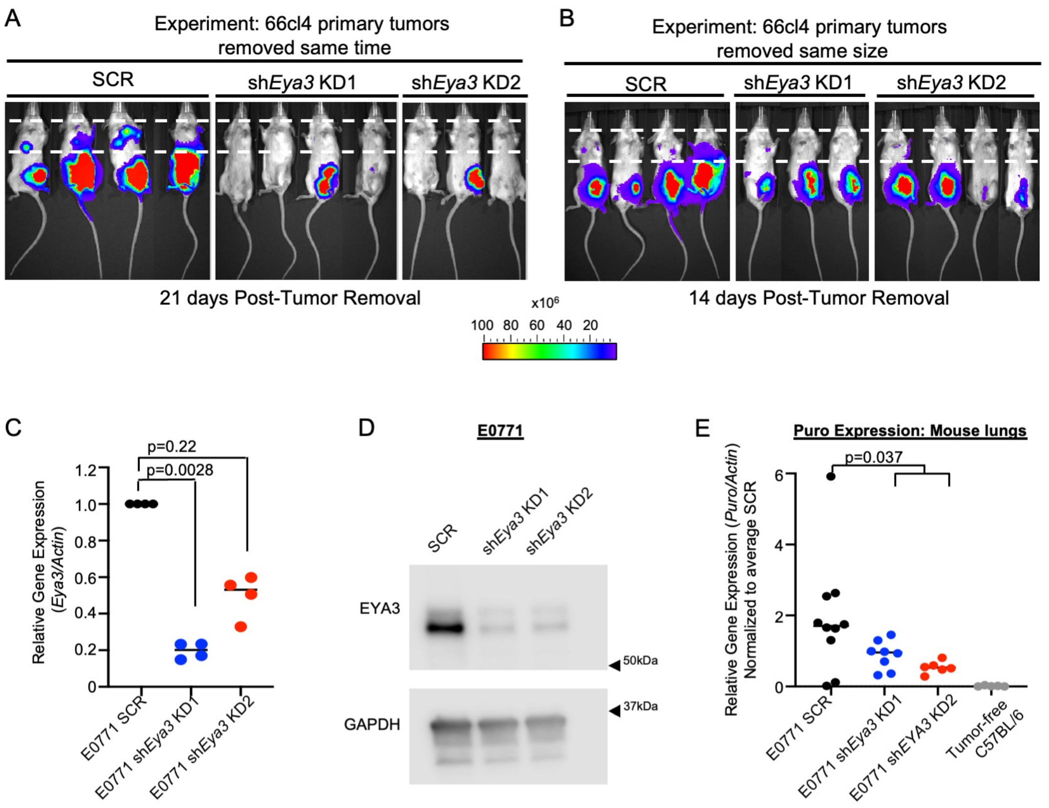


**Supplemental Figure 1: EYA3 regulates metastasis of 66cl4 and E0771 cells**
(A) Related to Figure 1F, bioluminescence was detected with the IVIS200. Shown are representative images of mice captured 21 days after primary tumor removal. (B) Related to Figure 1H, bioluminescence was detected with the IVIS200. Shown are representative images of mice captured 21 days after primary tumor removal. Scale for A and B shown below (units = photons/sec/$cm^2$). (C) *Eya3* expression determined by qRT-PCR in E0771 cells. (D) EYA3 protein levels as determined by Western blot analysis in E0771 cells. (E) 500,000 E0771 SCR, sh*Eya3* KD1, or sh*Eya3* KD2 were injected intravenously into C57BL/6 mice. Mice were sacrificed 21-23 days after intravenous injection of cancer cells and RNA was isolated from the lungs. Lung RNA was probed for expression of puromycin resistance gene by qRT-PCR and normalized to *Actin*, as a readout for metastatic cancer cells colonizing the lungs. *Puro/Actin* Ct values were averaged across all SCR samples, and *Puro/Actin* for each sample was normalized to this average value to determine fold change. RNA from the lungs of a tumor-free C57BL/6 mouse was used as a negative control. 1 mouse was excluded from KD1 due to degraded lung RNA sample, and 3 mice were excluded from KD2 due to premature death. ROUT (Q=5%) method identified one outlier in KD1, one outlier in KD2, and one outlier in the tumor-fee lung, which were removed. (SCR group, n=10; KD1 group, n=8; KD2 group, n=6). Statistical analysis was performed using ANOVA with sum contrasts in R.

Suppl. Figure 2

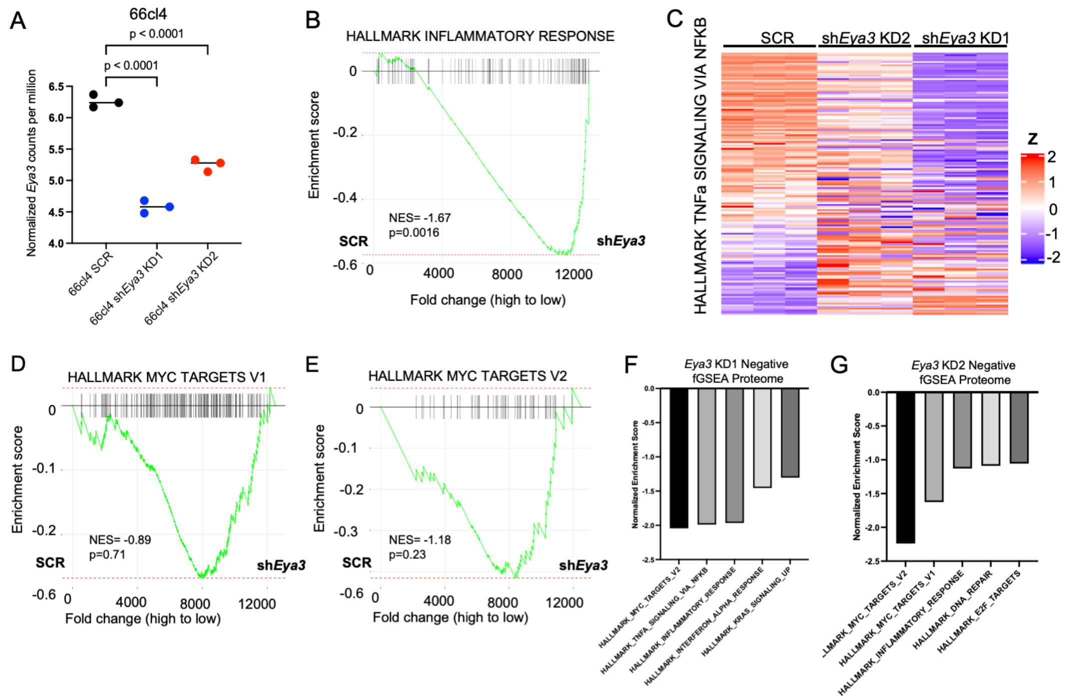


H

| | Motif | Transcription Factor | NES | # Enriched Genes |
|---|---|---|---|---|
| 1 | cisbp__M6511 | Rela (inferredBy_Orthology). | 5.4 | 61 |
| 2 | cisbp__M4496 | Rela (inferredBy_Orthology). | 5.3 | 33 |
| 3 | jaspar__MA0107.1 | Rela (inferredBy_Orthology). | 5.28 | 37 |
| 4 | hocomoco__TF65_MOUSE. H11MO.0.A | Rela (directAnnotation). | 5.11 | 49 |
| 5 | hocomoco__TF65_HUMAN. H11MO.0.A | Rela (inferredBy_Orthology). | 5.08 | 33 |
| 6 | cisbp__M6449 | Rel (inferredBy_Orthology). | 5.06 | 29 |
| 7 | transfac_public__M00052 | Rela (directAnnotation). | 4.84 | 39 |
| 8 | cisbp__M4468 | Rela (inferredBy_Orthology). | 4.82 | 39 |
| 9 | cisbp__M4498 | Rela (inferredBy_Orthology). | 4.78 | 38 |
| 10 | cisbp__M4491 | Rela (inferredBy_Orthology). | 4.77 | 24 |

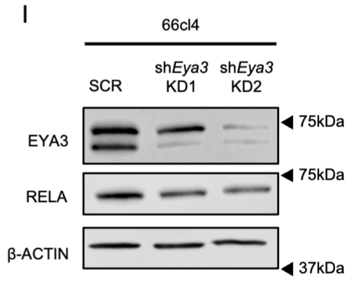


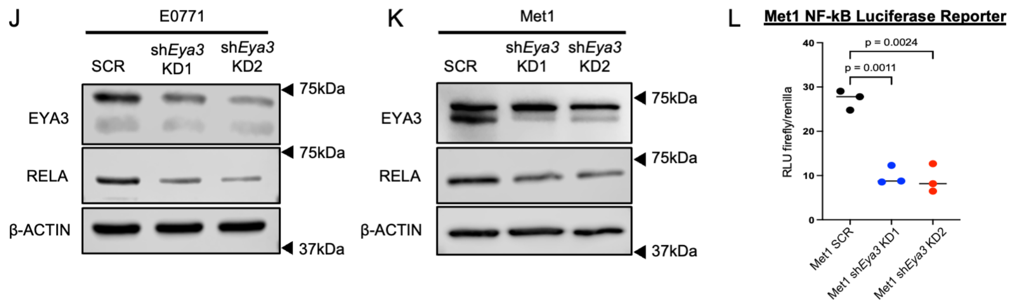


**Supplemental Figure 2: EYA3 regulates immune, NF-kB, and MYC gene signatures/transcriptional targets**

(A) *Eya3* mRNA levels (counts per million) in 66cl4 SCR, sh*Eya3* KD1, and sh*Eya3* KD2 cells used in RNA-sequencing experiment. (B) GSEA plot of Hallmark Inflammatory Response gene set between 66cl4 SCR and combined *Eya3* KD cells. (C) Heat map of DEGs in the Hallmark TNFa Signaling Via NFKB gene set from the RNA-sequencing. (D) GSEA plot for Hallmark Myc Targets V1 gene set for 66cl4 SCR vs combined sh*Eya3* KD1 and sh*Eya3* KD2 groups from aforementioned RNA-sequencing experiment. (E) GSEA plot for Hallmark Myc Targets V2 gene set for 66cl4 SCR vs combined sh*Eya3* KD1 and shEya3 KD2 groups from aforementioned RNA-sequencing experiment. (F) Top downregulated Hallmark gene sets (by normalized enrichment score) from proteome experiment for 66cl4 sh*Eya3* KD1 vs. SCR cells. (G) Top downregulated Hallmark gene sets (by normalized enrichment score) from proteome experiment for 66cl4 sh*Eya3* KD2 vs. SCR cells. (H) Top ten hits from transcription factor motif enrichment analysis of DEGs between 66cl4 SCR and *Eya3* KD cells (cutoff log2 FC>1). (I-K) Western blot analyses of EYA3 and RELA (B-ACTIN levels shown as a loading control) whole cell lysates in (I) 66cl4, (J) E0771, and (K) Met1 SCR and sh*Eya3* KD cells. (L) NF-kB luciferase reporter assay comparing transcriptional activation of NF-kB response element in Met1 SCR and *Eya3* KD cells.

Suppl. Figure 3

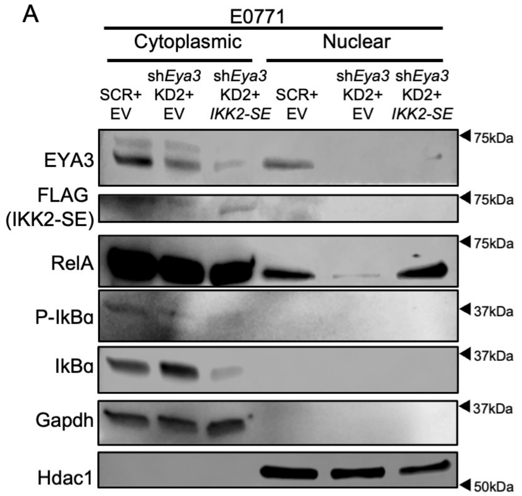


C

GSEA Analysis

sh*Eya3* KD2+*IKK2-SE* vs. sh*Eya3* KD2+EV

| Rank | Hallmark Gene Set | NES | P.Adj |
|---|---|---|---|
| 1 | INFLAMMATORY_RESPONSE | 2.22099795 | 0.00132679 |
| 2 | INTERFERON_GAMMA_RESPONSE | 2.18054554 | 0.00132679 |
| 3 | IL6_JAK_STAT3_SIGNALING | 2.08982755 | 0.00132679 |
| 4 | ALLOGRAFT_REJECTION | 2.08646702 | 0.00132679 |
| 5 | INTERFERON_ALPHA_RESPONSE | 2.05551115 | 0.00132679 |
| 6 | COMPLEMENT | 1.86305 | 0.00253233 |
| 7 | TNFA_SIGNALING_VIA_NFKB | 1.85194597 | 0.00188281 |
| 8 | KRAS_SIGNALING_UP | 1.73463133 | 0.02133931 |
| 9 | EPITHELIAL_MESENCHYMAL_TRANSITION | 1.62761921 | 0.05060075 |
| 10 | APOPTOSIS | 1.47586055 | 0.13074392 |
| 11 | ANGIOGENESIS | 1.44549403 | 0.24005196 |
| 12 | COAGULATION | 1.40921383 | 0.23257545 |
| 13 | P53_PATHWAY | 1.38465628 | 0.15683741 |
| 14 | XENOBIOTIC_METABOLISM | 1.38245253 | 0.20415306 |
| 15 | ESTROGEN_RESPONSE_EARLY | 1.25478416 | 0.34597221 |
| 16 | IL2_STAT5_SIGNALING | 1.2359585 | 0.36158586 |
| 17 | APICAL_SURFACE | 1.18461063 | 0.46696255 |
| 18 | KRAS_SIGNALING_DN | 1.174622 | 0.46696255 |
| 19 | APICAL_JUNCTION | 1.14582145 | 0.46696255 |
| 20 | REACTIVE_OXYGEN_SPECIES_PATHWAY | 1.10310916 | 0.5481109 |

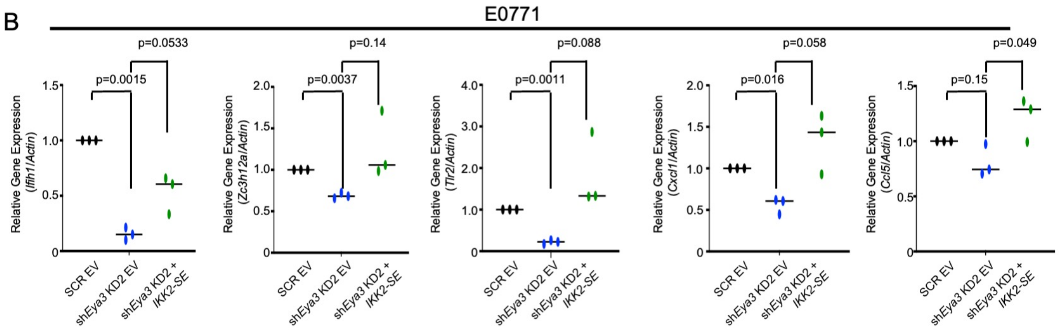


**Supplemental Figure 3: IKK2-SE rescues nuclear RELA levels in *Eya3* KD cells, restoring NF-kB gene expression but not MYC signaling**

(A) Western blot analysis of cytoplasmic and nuclear protein lysates from E0771 SCR+EV, sh*Eya3* KD2+EV, and sh*Eya3* KD2+IKK2-SE cells. GAPDH used as a cytoplasmic loading control. HDAC1 used as a nuclear loading control. (B) qRT-PCR analysis of NF-kB target genes *Ifih1, Zc3h12a, Tlr2, Cxcl1*, and *Ccl5* in E0771 SCR + EV, E0771 sh*Eya3* KD2 + EV, and E0771 shE*ya3* KD2 + *IKK2-SE* cells. Statistical analysis performed using Welch's unpaired t test. Mean±SD shown. (C) Top 20 Hallmark gene sets (as determined by normalized enrichment score [NES]) between 66cl4 sh*Eya3* KD2+*IKK2-SE* cells and 66cl4 sh*Eya3* KD2+EV cells.

Suppl. Figure 4

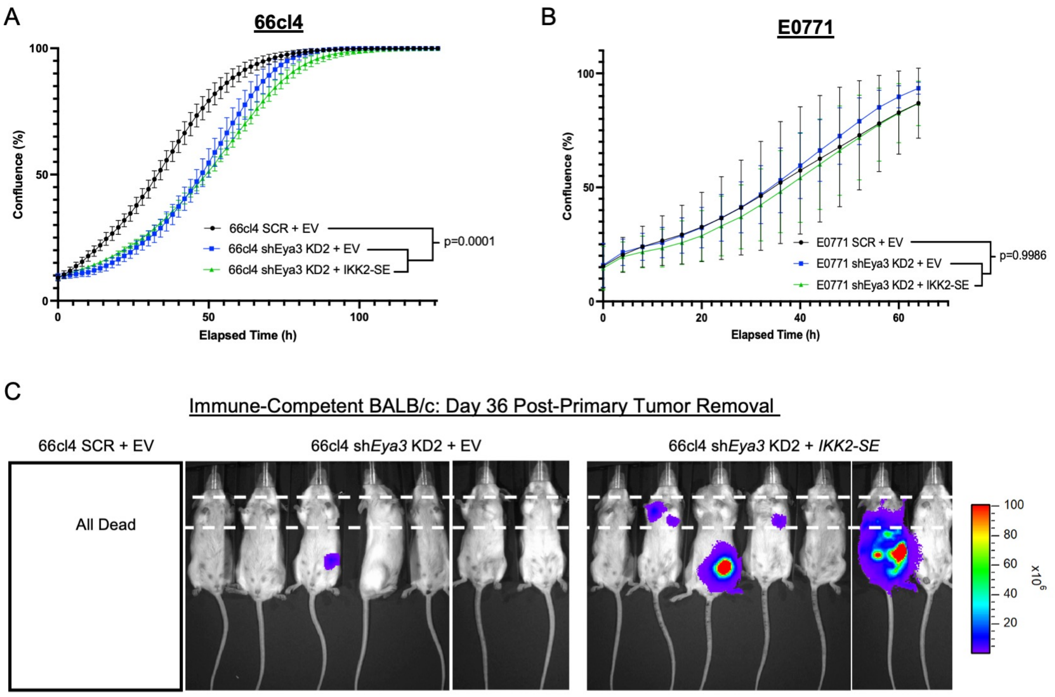


**Supplemental Figure 4: IKK2-SE does not rescue proliferation, but rescues lung metastasis, in *Eya3* KD TNBC cell lines**

(A-B) Incucyte analysis of cell confluence (technical replicates n = 5 per group) over time (hours) for SCR + EV, sh*Eya3* KD2 + EV, and shE*ya3* KD2 + *IKK2-SE* groups in the (A) 66cl4 and (B) E0771 cell lines. (C) Bioluminescent imaging of BALB/c immune-competent mice injected into the mammary fat pad with 66cl4 SCR + EV, sh*Eya3* KD2 + EV, and sh*Eya3* KD2 + *IKK2-SE* cells (n=7 mice/group) shown on Day 36 after primary tumor removal, related to Fig. 3E. Scale shown on right (units = photons).

Suppl. Figure 5

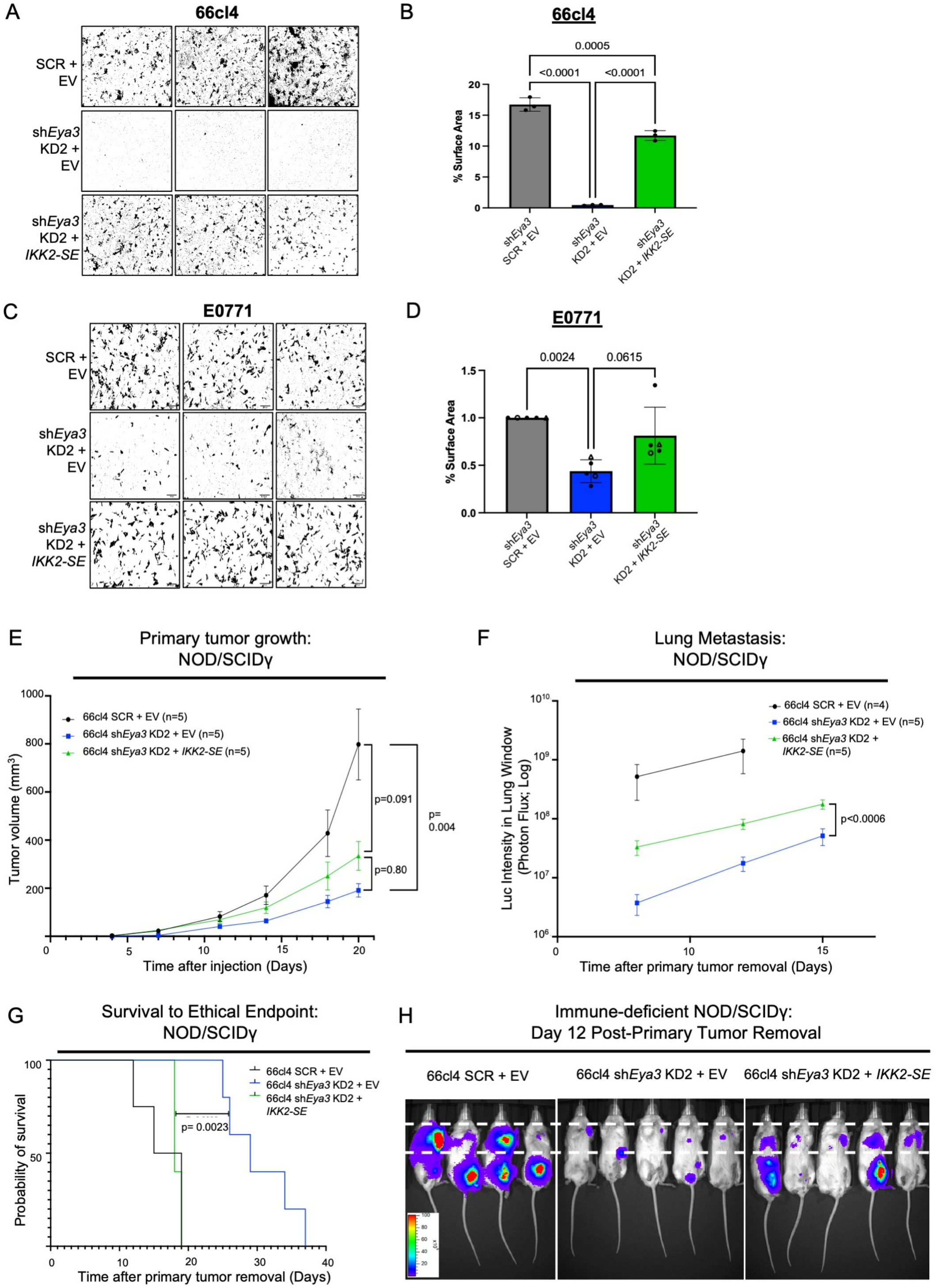


**Supplemental Figure 5: IKK2-SE does not rescue primary tumor growth, but modestly rescues metastasis in 66cl4 *Eya3* KD cells in NSG immune-deficient mice**

(A) Black and white mask of representative images from 24-hour transwell migration assays comparing 66cl4 SCR+EV, sh*Eya3* KD2+EV, and sh*Eya3* KD2+IKK2-SE cells. (B) Quantification of three independent replicates of the transwell migration assay shown in D. Statistical analysis performed using One-way ANOVA with post-hoc Tukey's multiple comparisons test. (C), As in A, for E0771. (D) Quantification of five independent replicates (at three time points: 8h = open circle, 10h = triangle, 12h = closed circle) of transwell migration assays shown in B. Statistical analysis performed using One-way ANOVA with post-hoc Tukeys multiple comparisons test. (E) Primary tumor growth, as measured by caliper measurement, of 66cl4 SCR+EV, sh*Eya3* KD2+EV, and sh*Eya3* KD2+*IKK-SE* cells after orthotopic injection into mammary fat pad of Nod-scid-gamma (NSG) mice. (F) Bioluminescence intensity in lung window post-tumor removal for NSG mice injected with 66cl4 SCR+EV, sh*Eya3* KD2+EV, and sh*Eya3* KD2 +*IKK2-SE* cells. Statistical analysis for A-B performed using longitudinal mixed model in R. (G) Overall survival to ethical endpoint for mice from B. Statistical analysis performed using log-rank test to compare survival. (H) Bioluminescent imaging of NSG immune-deficient mice injected into the mammary fat pad with 66cl4 SCR + EV, sh*Eya3* KD2 + EV, and sh*Eya3* KD2 + *IKK2-SE* cells (n=5 mice/group, one mouse died prior to this imaging time point in the SCR+EV group) shown on Day 12 after primary tumor removal. Bioluminescence was detected with the IVIS200. Scale shown on right (units = photons/sec/cm²).

Suppl. Figure 6

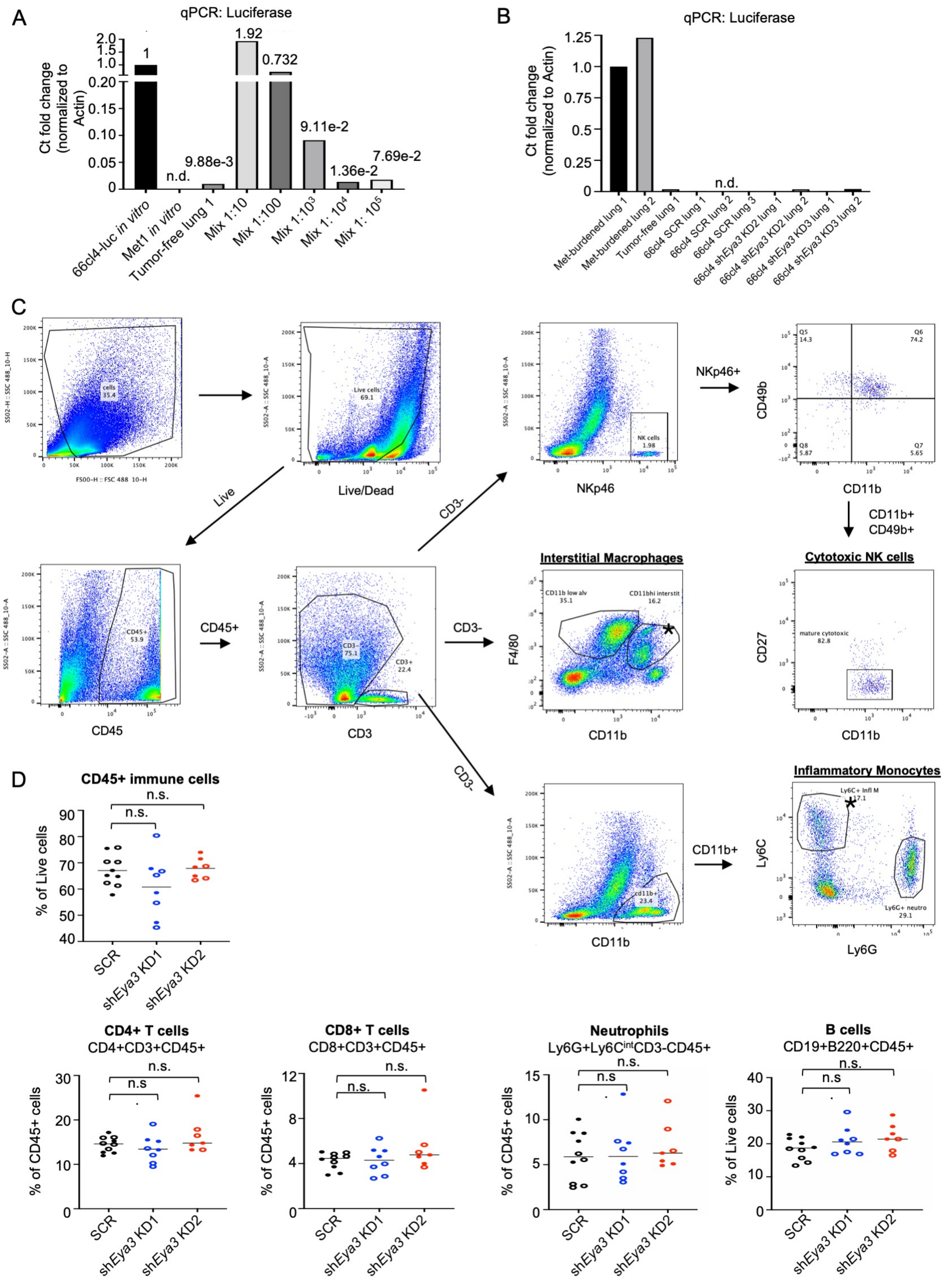


**Supplemental Figure 6: EYA3 regulates immune cell infiltration in the pre-metastatic lung, but has no effect on T cells, neutrophils, or B cells**

(A) RNA was isolated from cells *in vitro* that express luciferase (66cl4-luc) or do not express luciferase (Met1), as well as from the lung of a mouse that did not have any tumors or metastases for use as positive and negative controls. cDNA from 66cl4-luc was mixed with cDNA from the tumor-free lung at varying ratios and tested for luciferase by qPCR to determine the detection limit of the assay. (B) RNA was isolated from mouse lungs and analyzed for luciferase by qPCR to detect the presence of overt micrometastases. (C) Shown is the gating strategy for data related to Figs. 4, 5, and 7. Interstitial macrophages were considered as Zombie-CD45+CD3-F4/80$^{hi}$CD11b+; Inflammatory monocytes: Zombie-CD45+CD3-CD11b+Ly6C+Ly6C-; and Cytotoxic NK cells: Zombie-CD45+CD3-NKp46+CD49b+CD11b+CD27- in BALB/c and Zombie-CD45+CD3-NKp46+CD49b+CD11b+ in C57BL/6 (as in this model nearly all NK cells were uniformly dually positive for CD11b and CD27). (D) Related to Fig. 4, lung single cell suspensions from pre-metastatic lungs were analyzed for CD45+ lymphocytes, T cells, neutrophils, and B cells by flow cytometry. Statistical analysis was performed using Kruskal-Wallis or One-way ANOVA, adjusted for multiple comparisons.

Suppl. Figure 7

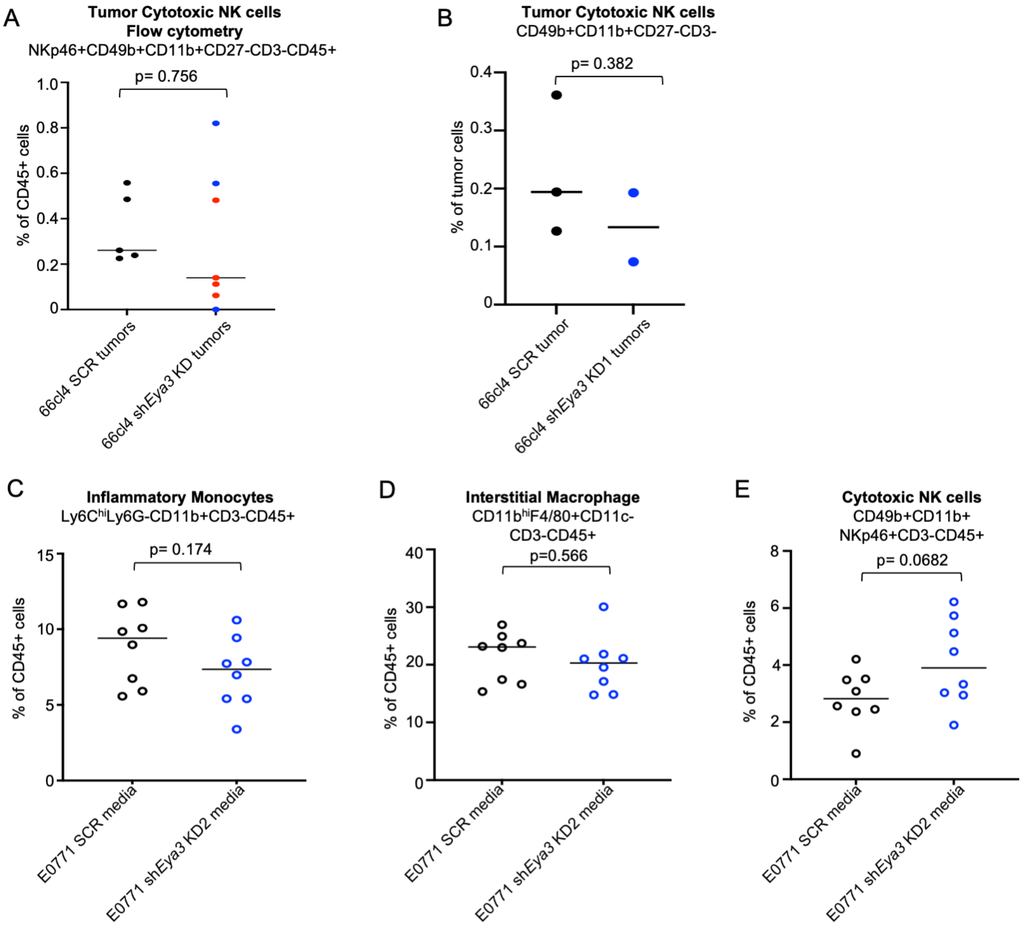


**Supplemental Figure 7. *Eya3* expression does not influence cytotoxic NK cell infiltration in the early primary tumor, but CM from EYA3 regulates cytotoxic NK cell infiltration to the pre-metastatic lung in multiple models**

(A) When tumor size exceeded 50mm$^3$, mice were sacrificed and primary tumor single cell suspensions were analyzed by flow cytometry for NK cells. sh*Eya3* KD1 and KD2 groups were combined due to tumor growth inhibitory effects of Eya3 and subsequent small sample size (n=7 total), with KD1 in blue and KD2 in red. (B) When tumor size exceeded 50mm$^3$, mice were sacrificed and primary tumors were fixed in formalin. SCR and sh*Eya3* KD1 tumor slices were stained with an NK-focused antibody panel, imaged on the Vectra Polaris, and scored for various cell populations using inForm image analysis software. Each dot represents a different tumor. Statistical analysis was performed using Welch's unpaired t test. (C-E) E0771 SCR, *shEya3* KD1, and *shEya3* KD2 cells were orthotopically injected into the fourth mammary fat pad of syngeneic C57BL/6 mice. When tumor size exceeded 50mm$^3$, mice were sacrificed. (C-E) Lung single cell suspensions were analyzed for (C) inflammatory monocytes, (D) interstitial macrophages, and (E) cytotoxic NK cells by flow cytometry according to the gating strategy detailed in Suppl. Fig. 5.

Suppl. Figure 8

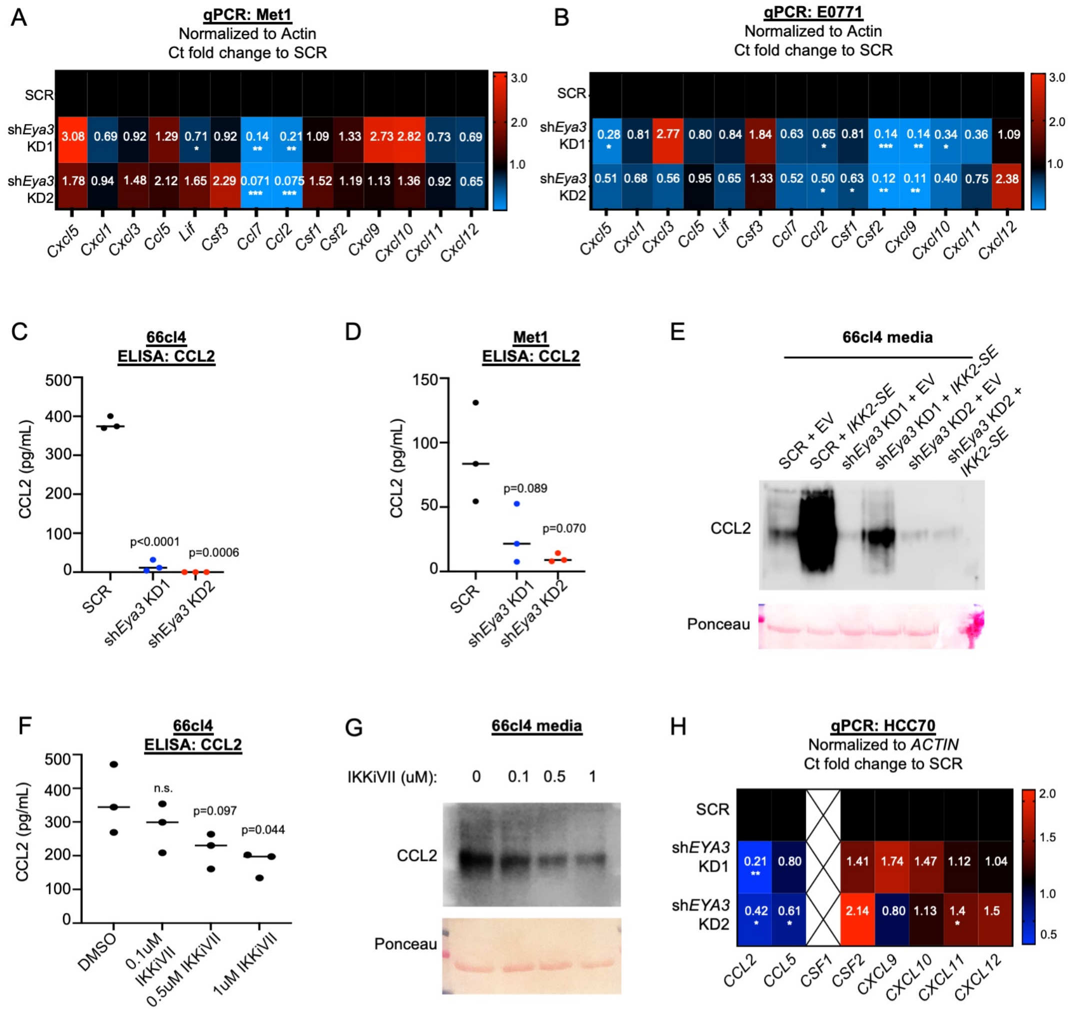


**Supplemental Figure 8. EYA3 regulates the secreted factor CCL2 downstream of NF-kB**
(A) Met1 cells with or without *Eya3* knockdown were probed for cytokine expression by qRT-PCR and normalized to Actin. Fold change was calculated compared to SCR cells, and data were averaged from three independent experiments. (B) As in A, for E0771 cells. Data were averaged from 3-5 independent experiments. (C-D) Media was incubated with 66cl4 (C) or Met1 (D) cells for 48 hours, and then probed for CCL2 by ELISA assay. Data represent an average of three independent experiments. (E) Media was incubated with 66cl4 cells with or without *Eya3* KD, with or without *IKK2-SE* expression for 48 hours. Conditioned media was then probed for CCL2 by Western blot. Shown is a representative image of three independent experiments. (F) 66cl4 cells were treated with varying concentrations of IKK inhibitor VII for 24 hours in serum-free media. Conditioned media was probed for CCL2 by ELISA assay. Data represent an average of three independent experiments. (G) Cells were treated as in F, and conditioned media was probed for CCL2 by Western blot. Shown is a representative image of three independent experiments. (H) As in A, for HCC70 cells.

Suppl. Figure 9

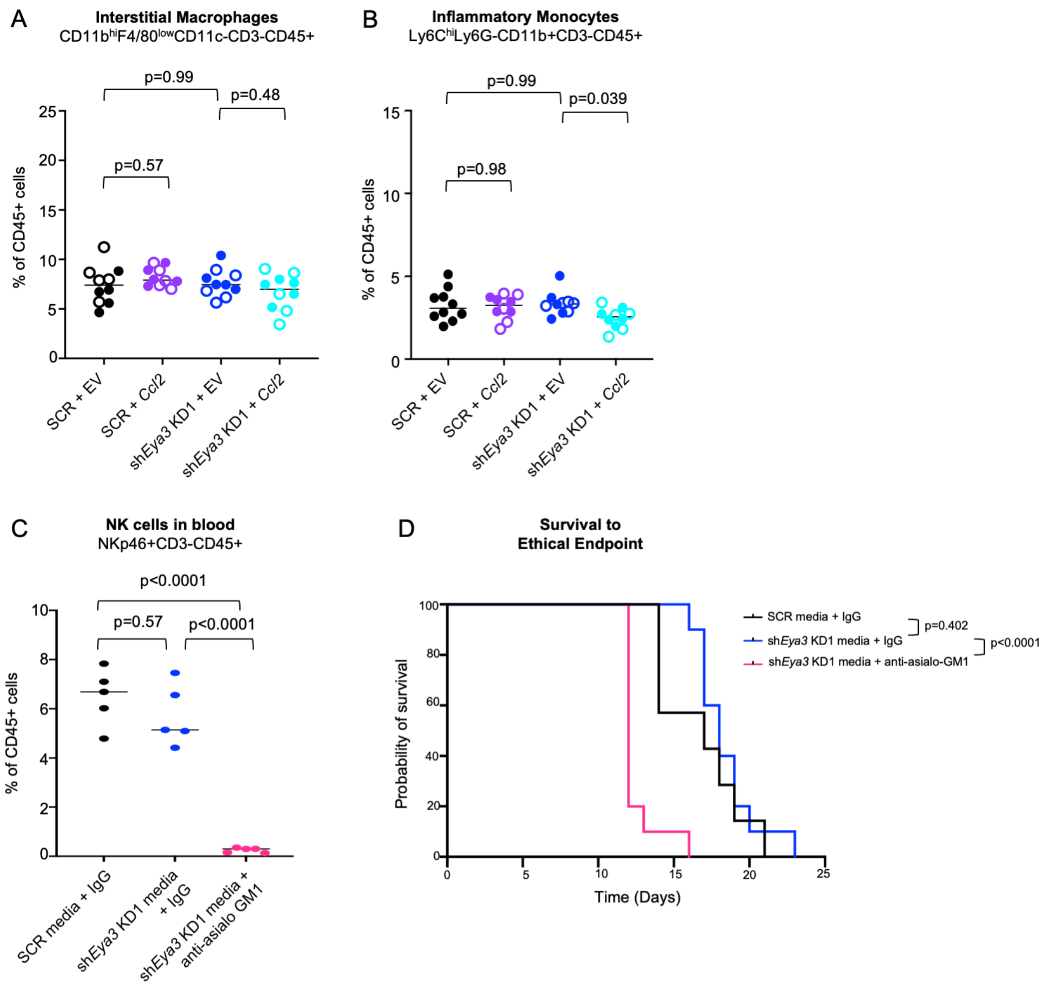


**Supplemental Figure 9. *Ccl2* re-expression in conditioned media does not alter infiltration of interstitial macrophages and monocytes to the lungs**
(A-B) Related to Fig. 7, interstitial macrophages (A) and inflammatory monocytes (B) were analyzed by flow cytometry in the lungs of mice treated with conditioned media. Data were collected from 10 mice per group, from two independent experiments. Statistical analysis was performed using Kruskal-Wallis or One-way ANOVA, adjusted for multiple comparisons. (C) NK cell depletion was confirmed on a subset of mice through flow cytometry staining of blood collected by submandibular bleeds one day prior to tumor cell injection using antibodies against NKp46, CD3 and CD45. (D) Overall survival to ethical endpoint of mice injected with 66cl4 SCR and treated with either SCR media + IgG (N=7), sh*Eya3* KD1 media + IgG (N=10), or sh*Eya3* KD1 media + anti-asialo GM1 (N=10) after tail vein injection. Statistical analysis performed using log-rank test between groups indicated.

Suppl. Figure 10

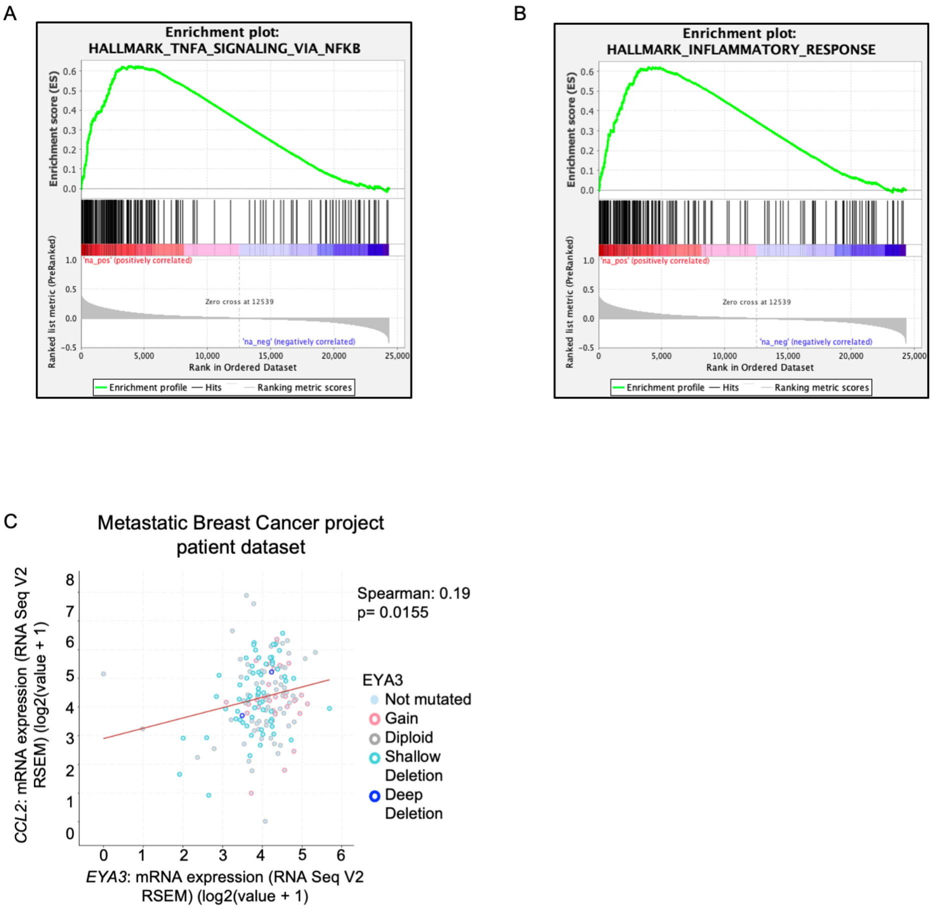


**Supplementary Figure 10. *EYA3*, NF-kB signaling, and *CCL2* levels are correlated in multiple patient datasets**

(A) GSEA plot of Hallmark TNFa signaling via NF-kB gene set correlated to *EYA3* expression from analysis in Fig. 8A. (B) GSEA plot of Hallmark Inflammatory Response gene set correlated to *EYA3* expression from analysis in Fig. 8A. (C) Correlation of *CCL2* and *EYA3* mRNA expression in the Metastatic Breast Cancer project patient dataset on cBioPortal.

**Supplemental Material and Methods**

qRT-PCR primers:

| | |
|---|---|
| Mouse Eya3 FWD | CAGAGCAGAGATCGAGGTGC |
| Mouse Eya3 REV | CTGGAACCAACTGCGTGGTA |
| Mouse B-actin FWD | AGGCATTGTGATGGACTCCG |
| Mouse B-actin REV | ATGTCACGCACGATTTCCCT |
| Mouse Gapdh FWD | TGTGAACGGATTTGGCCGTA |
| Mouse Gapdh REV | ACTGTGCCGTTGAATTTGCC |
| Mouse Rela FWD | CTTCCTCAGCCATGGTACCTCT |
| Mouse Rela REV | CAAGTCTTCATCAGCATAACTG |
| Mouse Relb FWD | CTTTGCCTATGATCCTTCTGC |
| Mouse Relb REV | GAGTCCAGTGATAGGGGCTCT |
| Mouse Nfkb1 FWD | GAAATTCCTGATCCAGACAAAAAC |
| Mouse Nfkb1 REV | ATCACTTCAATGGCCTCTGTGTAG |
| Mouse Nfkb2 FWD | CTGGTGGACACATACAGGAAGAC |
| Mouse Nfkb2 REV | ATAGGCACTGTCTTCTTTCACCTC |
| Mouse Pp2a-b55a FWD | GCCACAGGAGATAAAGGTGGG |
| Mouse Pp2a-b55a REV | ATTCTGGCTCATGGCTCTGG |
| Puromycin-FWD | TCACCGAGCTGCAAGAACTCT |
| Puromycin-REV | CCCACACCTTGCCGATGT |
| Mouse Ifih1-FWD | GTGATGACGAGGCCAGCAGTTG |
| Mouse Ifih1-REV | ATTCATCCGTTTCGTCCAGTTTCA |
| Mouse Zc3h12a-FWD | CTGCCTCTCAGTCCAGCTCT |
| Mouse Zc3h12a-REV | GGAGTGAGTCCTGGGTGTGT |
| Mouse Tlr2-FWD | GCCACCATTTCCACGGACT |
| Mouse Tlr2-REV | GGCTTCCTCTTGGCCTGG |
| Mouse Cxcl1-FWD | AAGAATGGTCGCGAGGCTTG |
| Mouse Cxcl1-REV | AGGTGCCATCAGAGCAGTCT |
| Mouse Ccl5-FWD | TGCCCACGTCAAGGAGTATT |

| | |
|---|---|
| Mouse Ccl5-REV | CAGGACCGGAGTGGGAGTA |
| Mouse Ccl2-FWD #1 | CCCAATGAGTAGGCTGGAGA |
| Mouse Ccl2-REV #1 | AAAATGGATCCACACCTTGC |
| Mouse Ccl2-FWD #2 | AGTAGGCTGGAGAGCTACAA |
| Mouse Ccl2-REV #2 | GTATGTCTGGACCCATTCCTTC |
| Mouse Csf1-FWD | CATCCAGGCAGAGACTGACA |
| Mouse Csf1-REV | CTTGCTGATCCTCCTTCCAG |
| Mouse Csf2-FWD | TGGAAGCATGTAGAGGCCATCA |
| Mouse Csf2-REV | GCGCCCTTGAGTTTGGTGAAAT |
| Mouse Cxcl12-FWD | GCACGGCTGAAGAACAACAAC |
| Mouse Cxcl12-REV | TTCCTCGGGCGTCTGACTC |
| Mouse Cxcl10-FWD | CCATCAGCACCATGAACCCAAGT |
| Mouse Cxcl10-REV | CACTCCAGTTAAGGAGCCCTTTTAGACC |
| Mouse Cxcl5-FWD | TGATCGCTAATTTGGAGGTGAT |
| Mouse Cxcl5-REV | TAGCTTTCTTTTTGTCA |
| Mouse Cxcl3-FWD | AGATCTCACCACAGCCCTTC |
| Mouse Cxcl3-REV | AACCCTTGGTAGGGTGTTCA |
| Mouse Lif-FWD | CTGCTGGTTCTGCACTGGAAAC |
| Mouse Lif-REV | GCTCCCCTTGAGCTGTGTAATAG |
| Mouse Csf3-FWD | TTGGCAACATCCAGCTGAAG |
| Mouse Csf3-REV | GCAGGCTCTATCGGGTATTTCC |
| Mouse Ccl7-FWD | TCAAGAGCTACAGAAGGATCACC |
| Mouse Ccl7-REV | TGGAGTTGGGGTTTTCATGTCT |
| Human CCL2-FWD #1 | CCCCAGTCACCTGCTGTTAT |
| Human CCL2-REV #1 | TGGAATCCTGAACCCACTTC |
| Human CCL2-FWD #2 | GCTCAGCCAGATGCAATCAA |
| Human CCL2-REV #2 | TTCTTTGGGACACTTGCTGC |
| Human CCL5-FWD | ACACCCTGCTGCTTTGCCTACA |
| Human CCL5-REV | TCCCGAACCCATTTCTTCTCTG |

| | |
|---|---|
| Human CXCL9-FWD | GGTGTTCTTTTCCTCTTGGGC |
| Human CXCL9-REV | AACAGCGACCCTTTCTCACT |
| Human CXCL10-FWD | CCCACGTGTTGAGATCATTG |
| Human CXCL10-REV | TCCATCACAGCACCGGG |
| Human CXCL11-FWD | GAGTGTGAAGGGCATGGCTA |
| Human CXCL11-REV | ATGCAAAGACAGCGTCCTCT |
| Human CXCL12-FWD | TCAGCCTGAGCTACAGATGC |
| Human CXCL12-REV | CTTTAGCTTCGGGTCAATGC |
| Human CSF2-FWD | GGCGTCTCCTGAACCTGAGT |
| Human CSF2-REV | GGGGATGACAAGCAGAAAGT |
| Luciferase-FWD | GGGATACGACAAGGATATGGGC |
| Luciferase-REV | TGGAACAACTTTACCGACCGC |

Western blot antibodies (primary):

| **Reagent – Primary Antibodies** | **Source** | **Identifier** | **Clone number (if applicable)** | **Dilution used** |
|---|---|---|---|---|
| Rabbit polyclonal anti-Eya3 | Bethyl Laboratories | A302-689A | | 1:500 |
| Rabbit polyclonal anti-RelA (p65) | Sigma-Aldrich | 06-418 | | 1:500 |
| Rabbit monoclonal anti-Gapdh | Cell Signaling Technology | 5174S | D16H11 | 1:1000 |
| Mouse monoclonal anti-β-Actin | Sigma-Aldrich | A5316 | AC-74 | 1:3000 |
| Mouse monoclonal anti-HDAC1 | Santa Cruz Biotechnology | Sc-81598 | 10E2 | 1:100 |
| Mouse monoclonal anti-p-IkBa (Ser32/Ser36) | Cell Signaling Technology | 9246S | 5A5 | 1:1000 |
| Rabbit polyclonal anti-IkBa | Cell Signaling Technology | 9242S | | 1:1000 |
| Mouse monoclonal anti-FLAG | Sigma-Aldrich | F1804 | Clone M2 | 1:1000 |
| Mouse CCL2 (MCP-1) | Cell Signaling | 2029 | | 1:1000 |

Western blot antibodies (secondary):

| Reagent – Secondary Antibody | Source | Identifier | Dilution Used |
|---|---|---|---|
| HRP-conjugated goat anti-mouse | Li-Cor | 926-80010 | 1:10000 |
| HRP-conjugated goat anti-rabbit | Li-Cor | 926-80011 | 1:10000 |

Flow cytometry antibodies:

| Reagent – Primary Antibodies | Source | Clone | Dilution Used |
|---|---|---|---|
| CD45 | BioLegend | 30-F11 | 1:400 |
| CD3 | BioLegend | 17A2 | 1:200 |
| F4/80 | BioLegend | BM8 | 1:400 |
| Ly6C | BioLegend | HK1.4 | 1:1600 |
| Ly6G | BioLegend | 1A8 | 1:400 |
| CD11b | eBiosciences | M1/70 | 1:200 |
| CD11c | BioLegend | N418 | 1:200 |
| MHC-II (I-A/I-E) | BioLegend | M5/114.15.2 | 1:1600 |
| CD206 | BioLegend | C068C2 | 1:200 |
| CD80 | Thermo Fisher Scientific | 16-10A1 | 1:300 |
| PD-L1 | BioLegend | 10F.9G2 | 1:400 |
| NKp46 | BioLegend | 29A1.4 | 1:133 |
| CD49b | BioLegend | DX5 | 1:133 |
| CD27 | BioLegend | LG.3A10 | 1:200 |
| CD8a | BioLegend | 53-6.7 | 1:400 |
| CD4 | eBiosciences | GK1.5 | 1:400 |
| Foxp3 | eBiosciences | FJK-16s | 1:400 |
| CD19 | BioLegend | 6D5 | 1:133 |
| B220 | BioLegend | RA3-6B2 | 1:400 |
| IFNɣ | BioLegend | XMG1.2 | 1:200 |
| TNFɑ | BioLegend | MP6-XT22 | 1:200 |

Vectra Staining

We injected 66cl4 cells with or without Eya3 knockdown into the mammary fat pad of syngeneic mice and allowed palpable tumors (>~50mm3) to form. Once tumors were palpable, mice were sacrificed and tumors were fixed in formalin. SCR and shEYA3 KD1 tumor slices were stained with an NK-focused antibody panel by the Human Immune Monitoring Shared Resource at

CU/AMC, imaged on the Akoya Biosciences' Vectra Polaris, and scored for various cell populations using inForm image analysis software.

Proteomics Sample Preparation, TMT labeling, and Phosphoenrichment

Cells were washed with PBS and immediately harvested with 5% (w/v) sodium dodecyl sulfate (SDS), 10 mM tris(2-carboxyethylphosphine) (TCEP), 40 mM 2-chloroacetamide and 50 mM Tris-HCl, pH 8.5 and boiled 10 minutes. Cell lysates were probe sonicated and stored at -80°C. Extracted proteins were digested using the SP3 method114. Briefly, 500 µg carboxylate-functionalized speedbeads (Cytiva Life Sciences) were added followed by the addition of acetonitrile to 80% (v/v) inducing binding to the beads. The beads were washed twice with 80% (v/v) ethanol and twice with 100% acetonitrile. Proteins were digested in 50 mM Tris-HCl, pH 8.5, with Lys-C/Trypsin (Promega) incubating at 37˚C shaking at 1000rpm overnight. Tryptic peptides were desalted using Waters HLB Oasis cartridge according to the manufacturer's instructions and dried in a speedvac vacuum centrifuge and stored at -20°C. Peptides were labeled with the Tandem Mass Tags (TMT) 10-plex (Thermo Scientific) according to the manufacturer's instructions. Briefly, peptides were suspended in 0.1M triethylammonium bicarbonate and each TMT label was added in acetonitrile and incubated for 1h at ambient. TMT Labeling reactions were quenched with the addition of hydroxylamine and incubated for 15 minutes at ambient then combined. The multiplexed samples were desalted using a Waters HLB Oasis cartridge according to the manufacturer's instructions, dried in a speedvac vacuum centrifuge and stored at -20°C. A small portion of the multiplexed samples was set aside for proteome analyses and the remaining was phosphoenriched using the High-Select TiO2 phosphoenrichment kit (Thermo Scientific). Samples were suspended in the Binding/Equilibration buffer, loaded onto the TiO2 column, washed and eluted. The elution and the unretained flow through fractions were dried immediately in a speedvac vacuum centrifuge. The TiO2 elution was stored at -20°C, while the dried unretained flow through fraction was subjected to a sequential phosphoenrichment with the High-Select Fe-NTA phosphoenrichment kit (Thermo Scientific). The dried TiO2 unretained fraction was suspended in the Fe-NTA Binding/Wash Buffer, loaded onto the Fe-NTA column, washed and eluted. The TiO2 and the Fe-NTA elutions were combined and dried in a speedvac vacuum centrifuge. The proteome and phosphoenriched fractions were fractionated on a rpC18 column with high pH mobile phases (0.1% ammonium hydroxide) using a Waters M-class UPLC with a PDA detector and a custom fabricated 0.5 mm X 150 mm UChrom rpC18 1.8 µm 120 Å (nanolcms) column with a gradient from 2% to 40% ACN in 80 minutes for the proteome and from 2% to 25% ACN in 50 minutes for the phosphoproteome. Fractions were concatenated for a total of 24 proteome fractions and 12 phosphoproteome fractions. High pH fractions were immediately dried in a speedvac vacuum centrifuge and stored at -20°C until LC/MS analysis.

Mass Spectrometry and Data Analysis

High pH fractionated TMT-labeled peptides were suspended in 3% (v/v) acetonitrile (ACN), 0.1% (v/v) trifluoroacetic acid (TFA) and directly injected onto a reversed-phase CSH rpC18 1.7 µm, 130 Å, 75 mm X 250 mm M-class column (Waters), using an Ultimate 3000 nanoUPLC (Thermo Scientific). For the proteome analyses, peptides were eluted at 300 nL/minute with a gradient from 2% to 20% ACN, 0.1% (v/v) formic acid in 40 minutes then to 40% ACN in 5 minutes and detected using a Q-Exactive HF-X mass spectrometer (Thermo Scientific). For the phosphoproteome analyses, peptides were eluted at 300 nL/minute with a gradient from 2% to 20% ACN, 0.1% (v/v) formic acid in 120 minutes then to 40% ACN in 5 minutes and detected using a Q-Exactive HF-X mass spectrometer (Thermo Scientific). Precursor mass spectra (MS1) were acquired at a resolution of 120,000 from 380 to 1580 m/z with an automatic gain control (AGC) target of 3E6 and a maximum injection time of 50 milliseconds. Precursor peptide ion isolation width for MS2 fragment scans was 0.7 m/z with a 0.2 m/z isolation offset, and the top 12 most intense ions were sequenced. All MS2 spectra were acquired at a resolution of

60,000 with higher energy collision dissociation (HCD) at 30% normalized collision energy. An AGC target of 1E5 and 100 milliseconds maximum injection time was used. Dynamic exclusion was set for 25 seconds with a mass tolerance of ±10 ppm. Rawfiles were searched against the Uniprot Mus musculus database UP000000589 downloaded 11/13/2020 using MaxQuant v1.6.14.0. Cysteine carbamidomethylation and the TMT label were considered fixed modifications, while methionine oxidation, protein N-terminal acetylation and phosphorylation at serine, threonine or tyrosine were searched as variable modifications. All peptide and protein identifications were thresholded at a 1% false discovery rate (FDR). Proteome and phosphoproteome data were normalized using the cyclic loess method, then differential protein and phospho-occupancy were calculated using Limma (Bioconductor.org) to generate a linear model for fold-change estimation and standard error then an Empirical Bayes implementation of an independent t-test where the null distribution is inferred from the global data, and treatment distribution is Bayesian updated. FDR (q-values) were then generated using the Benjamini-Hochberg limma interpretation.

NF-kB Luciferase Reporter Assays

NF-kB 3kB Firefly luciferase expression and a 3kB mutant plasmid were obtained as a generous gift from Rebecca Schweppe (originally obtained from Addgene 26699). For all lines tested (Met1 SCR, shEya3 KD1, and shEya2 KD2; Met1 SCR, shB55a KD1, and shB55a KD2) 20,000 cells/well were plated in 96 well plates and transfected 1μg/well of either 3kB reporter (5 wells/group) or 3kB mutant (3 wells/group), and 500ng/well of Renilla control construct into indicated wells with Lipofectamine 2000 transfection reagent (Invitrogen, 11-668-019). After transfection, plate was incubated O/N at 37□C. After 24h incubation, cells were stimulated with 100ng/ml of TNFa for 4 hours in incubator followed by cell lysis using the Dual-Luciferase Reporter Assay System (Promega E1910) following manufacturer's instructions. The luciferase intensity per well and relative intensity of Firefly:Renilla luciferase was measured and calculated using the Turner Biosystems Modulus microplate reader. Statistical analysis was performed using a One-way ANOVA with post-hoc Dunnett's multiple comparisons test.

IncuCyte cell growth assays

Cell growth was measured using IncuCyte Zoom (Essen Biosciences) Live-Cell Analysis Platform. 5,000 66cl4 SCR+EV, 66cl4 shEya3 KD2+EV, and 66cl4 shEya3 KD2+IKK2-SE cells were plated in 5 replicates in a 96-well plate and were imaged every 2 hours with a 4x objective. The percent confluence of each well at each time point was calculated using IncuCyte Zoom image processing software. All 5 replicates per group were averaged to provide a single average confluence per time point, and each group was normalized to a consistent confluence percentage (as defined as being within 1% point) set to time point 0h for all groups. Cell growth curves were generated by graphing average confluence and standard deviation across technical replicates for each group for each time point. Statistical differences between groups were calculated using a longitudinal mixed effects model that compares groups over repeated measures.

Cell migration assays

For transwell cell migration assays, 66cl4 SCR+EV, shEya3 KD2+EV, and shEya3 KD2+IKK2-SE cells were resuspended in 200ul/insert of serum-free media and transferred into cell culture inserts with 8μm pores (BD Falcon, 353097) (200,000 cells/triplicate inserts per group). Cell culture inserts were placed in 24-well companion dishes (Corning, 353504) containing 800μl of full serum media per well and put in the incubator at 37°C (5% $CO_2$) for 24 hours. After incubation, media inside of the insert was aspirated, and the bottom of each insert was fixed in 4% PFA for 10 minutes at RT. After fixation, the bottom of each insert was stained with 0.1% crystal violet solution for 45 minutes at RT and then washed with $ddH_2O$ to remove excess stain. Inserts were dried at RT for 24 hours prior to imaging on an Olympus CKX41 microscope.

Three representative images of each insert were taken at 10X magnification (imager was blinded to the treatment group of each insert). Images were analyzed and quantified using ImageJ.
